# Differential interactions of resting, activated, and desensitized states of the α7 nicotinic acetylcholine receptor with lipidic modulators

**DOI:** 10.1101/2022.04.08.487470

**Authors:** Yuxuan Zhuang, Colleen M Noviello, Ryan E Hibbs, Rebecca J Howard, Erik Lindahl

**Affiliations:** Department of Biochemistry and Biophysics, Science for Life Laboratory, Stockholm University, Solna, Sweden; Department of Neuroscience, University of Texas Southwestern Medical Center, Dallas, TX, USA; Department of Applied Physics, Swedish e-Science Research Center, KTH Royal Institute of Technology, Solna, Sweden

## Abstract

The α7 nicotinic acetylcholine receptor is a pentameric ligand-gated ion channel that modulates neuronal excitability, largely by allowing Ca^2+^ permeation. Agonist binding promotes transition from a resting state to an activated state, and then rapidly to a desensitized state. Recently, cryo-EM structures of the human α7 receptor in nanodiscs were reported in multiple conformations. These were selectively stabilized by inhibitory, activating, or potentiating compounds. However, the functional annotation of these structures, and their differential interactions with unresolved lipids and ligands, remain incomplete. Here, we characterized their ion permeation, membrane interactions, and ligand binding using computational electrophysiology, free-energy calculations, and coarse-grained molecular dynamics. In contrast to non-conductive structures in apparent resting and desensitized states, the structure determined in the presence of the potentiator PNU-120596 was consistent with an activated state permeable to Ca^2+^. Transition to this state was associated with compression and rearrangement of the membrane, particularly in the vicinity of the peripheral MX helix. An intersubunit transmembrane site was implicated in selective binding of either PNU-120596 in the activated state, or cholesterol in the desensitized state. This substantiates functional assignment of all three lipid-embedded α7-receptor structures with ion permeation simulations. It also proposes testable models of their state-dependent interactions with lipophilic ligands, including a mechanism for allosteric modulation at the transmembrane subunit interface.

## Introduction

Pentameric ligand-gated ion channels (pLGICs) are key mediators of electrochemical signal transduction in the nervous system (Lester et al., 2004). Upon the binding of neurotransmitters, the corresponding pLGICs open, allowing either anions or cations to cross the membrane for further signal transduction (Lynagh and Pless, 2014). The α7 nicotinic acetylcholine receptor (α7 nAChR), a subtype of the nicotinic superfamily of pLGICs, consists of five identical α7 subunits. It is an important part of the cholinergic nervous system; defects in this receptor are associated with neurological conditions including schizophrenia, Alzheimer′s disease, and autism spectrum disorders (Antonio-Tolentino and Hopkins, 2020; Bouzat et al., 2018; Uteshev, 2014). The α7 nAChR is also widely expressed in the immune system and is linked to inflammatory disease (Kalkman and Feuerbach, 2016; Zdanowski et al., 2015). Accordingly, this channel constitutes an important therapeutic target. Although no drugs in current practice specifically target the α7 nAChR (Bouzat et al., 2018), the type-II positive allosteric modulator PNU-120596 (PNU) has proved a valuable pharmacological tool (Andersen et al., 2016; Gulsevin et al., 2019; Hurst, 2005). It prolongs channel opening and creates long-lived burst clusters in functional recordings. Binding sites for PNU have been proposed by both mutagenesis studies (daCosta et al., 2011; Szabo et al., 2014), molecular docking to homology models (Newcombe et al., 2018; Young et al., 2008), and were recently resolved in a detergent-embedded structure (Zhao et al., 2021). However, we still lack comprehensive integration of functional, structural, and simulation data that explicitly links state specific binding of PNU and other modulators to lipid interactions and channel conduction.

With the help of cryogenic electron microscopy (cryo-EM), three structures of the human α7 nAChR in lipid nanodiscs were recently reported in conditions expected to inhibit (bound to the antagonist α-bungarotoxin), activate (bound to the agonist epibatidine and modulator PNU), or desensitize (bound to only epibatidine) the receptor (Noviello et al., 2021) (Figure 1A). As for most pLGICs (Nemecz et al., 2016), the α7 nAChR can be dissected into an agonist-binding extracellular domain (ECD), pore-forming transmembrane domain (TMD), and semi-disordered intracellular domain (ICD). All three domains are implicated in influencing ion permeation and selectivity (Gharpure et al., 2019; Noviello et al., 2021). The partially resolved ICD forms lateral portals thought to pass ions to the intracellular space, rather than the central vestibule (Figure 1B) (Gharpure et al., 2019). Each subunit TMD consists of four membrane-spanning helices (M1–M4) with the ring of M2 helices lining the central ion pathway of the pentamer (Figure 1B, 1D). The ECD contains a constriction at residue E97 (Figure 1E) and has a Ca^2+^ binding site (Galzi et al., 1996; Noviello et al., 2021) alongside the permeation pathway near the ECD-TMD junction.

**Figure 1.**
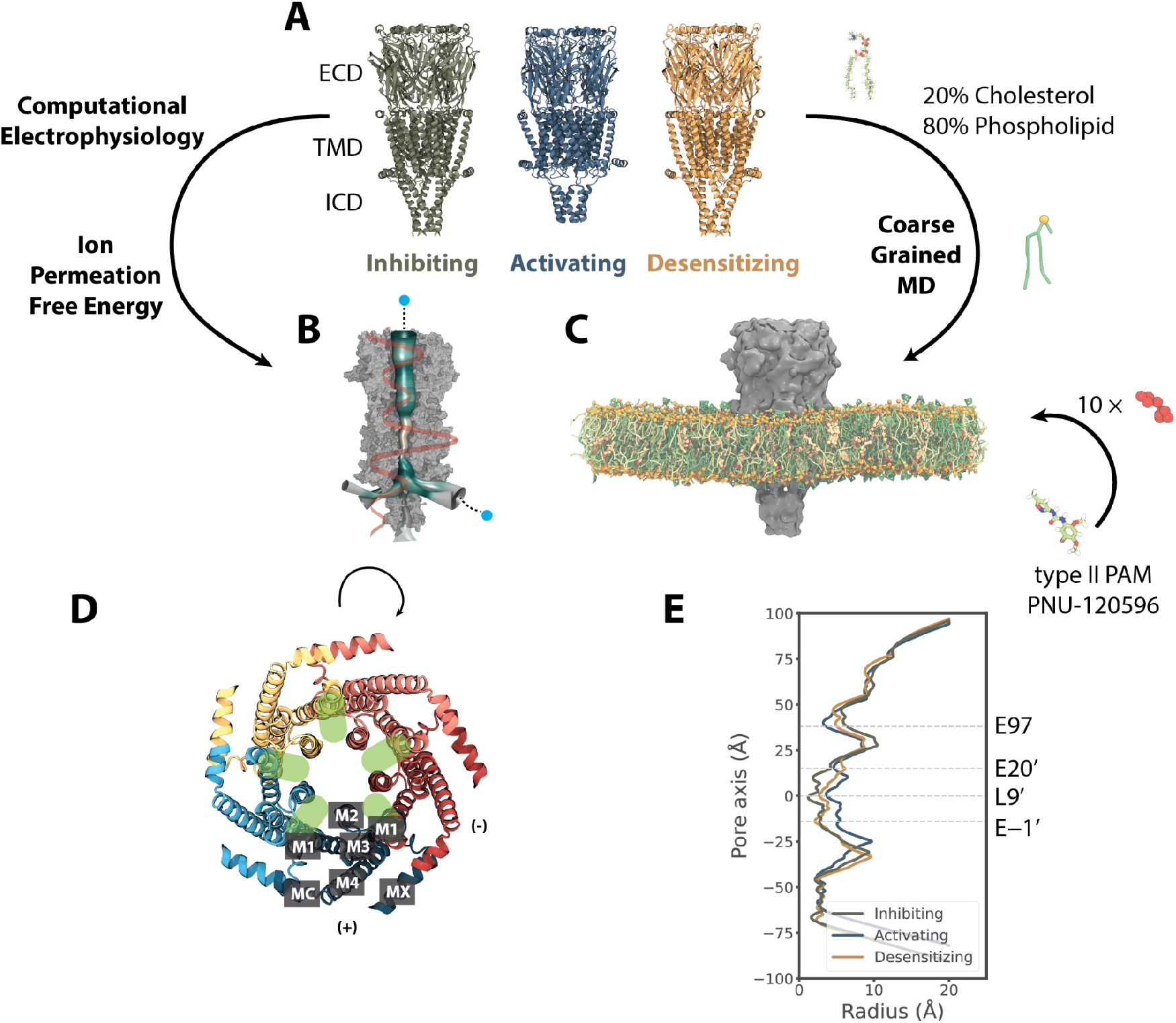
Computational approaches to test functional states and discover lipidic interactions of the human α7 nAChR. **A**. Cryo-EM structures of the α7 nAChR in lipid nanodiscs, viewed from the membrane plane, determined in inhibiting (olive, PDB ID 7KOO), activating (steel, PDB ID 7KOX), and desensitizing (orange, PDB ID 7KOQ) conditions, denoted as resting, activated, and desensitized states in this work, respectively. **B**. Computational electrophysiology and ion permeation free-energy profiles (orange curve) enable modelling of ion conductance and permeation. Mesh representation of the pore colored by hydrophobicity was generated by CHAP (Rao et al., 2019). **C**. Coarse-grained simulations enable quantification of protein interactions with mixed-lipid membranes (80% phospholipids, green; 20% cholesterol, yellow), as well as the ligand PNU (red). **D**. Details of the α7-nAChR TMD, showing the structure under activating conditions (PDB ID 7KOX) viewed from the extracellular side, colored by subunit. Labels on the bottom subunit indicate individual TMD helices M1–M4, MX, and MA. Green ovals indicate intersubunit cavities proximal to the channel pore. **E**. Pore-radius profiles (Rao et al., 2019) for the three experimental structures, colored as in A, with the midpoint (0 Å) of the channel axis at the 9′ hydrophobic gate. Dashed lines indicate key acidic residues, as well as the 9′ hydrophobic gate, facing the channel axis.

The α7 lipid-nanodisc structure obtained under activating conditions is strikingly different from those in either inhibiting or desensitizing conditions. In particular, the TMD helices are tilted and translated (Figure 1A), the surrounding membrane appears to be compressed, and the M4-MA helix partially loses its secondary structure. It remains to be determined whether this unanticipated state could account for distinctive permeation properties of the α7 subtype, e.g. relative selectivity for Ca^2+^ ions (Castro and Albuquerque, 1995; Noviello et al., 2021) and fast desensitization (Bouzat et al., 2008). Furthermore, comparing α7-nAChR structures highlights potentially important state-dependent interactions with the membrane. Membrane components including cholesterol have been shown to modulate nAChRs by binding directly to the TMD, as well as altering bulk lipid properties (Barrantes, 2004, 2012; John E.Baenziger, Jaimee A.Domville, J.P. Daniel Therien, 2017). Indeed, specific lipid interactions may critically influence the function of several pLGICs, especially in eukaryotes (Rosenhouse-Dantsker et al., 2012). In the case of α7, no specific lipids were resolved, leaving open questions as to the molecular details of state-specific remodeling at the protein-lipid interface.

Although the presumed activating conditions included a saturating concentration of PNU, this modulator could not be confidently resolved anywhere around the protein from the electron density. During preparation of this manuscript, a second set (Zhao et al., 2021) of α7-nAChR structures were reported under presumed resting (apo), partially desensitizing (agonist EVP-612 + PNU bound), and desensitizing (only EVP-612 bound) conditions. Although these were determined in detergent micelles, the structures in resting and desensitizing conditions were notably similar to those previously reported in lipid nanodiscs in inhibiting and desensitizing conditions, respectively. The PNU-bound structures were more divergent, with the detergent complex containing a narrower pore, less likely to represent a fully activated state. Moreover, PNU densities resolved in the intersubunit cavities of the detergent structure could not be superimposed in the lipid-nanodisc structure without steric clashes. Thus, the location and mechanism of PNU potentiation in the activated state remain unclear.

Molecular dynamics (MD) simulations complement experimental tools and have made it possible to study both ion permeation (Roux et al., 2004) and lipid/drug interactions of ion channels (Duncan et al., 2020; Hedger and Sansom, 2016). Here we report the first (to the best of our knowledge) MD study of three lipid-embedded experimental structures of the α7 nAChR, covering three key functional states. We tested the permeation properties of the structure under activating conditions utilizing both computational electrophysiology and ion-permeation calculations in comparison to experimental conductance and selectivity. We then quantified interactions of lipid molecules and PNU with different states of the channel, using coarse-grained simulations that make it possible to reach timescales where lipids diffuse to preferentially interact with different parts of the membrane protein. Our results detail lipidic interactions of specific residues in multiple TMD regions, and substantiate a mechanism of PNU-cholesterol dynamics that may underlie modulation of the α7-nAChR gating cycle.

## Results

### An open functional state under activating conditions

To test the functional state of the α7-nAChR structure determined under activating conditions, we first examined its ion conductance using computational electrophysiology (Kutzner et al., 2016, 2011). Two copies of the cryo-EM structure (PDB ID 7KOX, determined in the presence of epibatidine and PNU) were embedded in lipid bilayers oriented opposite one another in a single simulation box, enabling the generation of a range of electrostatic potentials in a ∼150-mM NaCl medium (Figure 2A). In the presence of voltage differences from -700 mV to 400 mV (Figure 2B), Na^+^ preferably permeated the channel, with a selectivity of ∼10:1 over Cl^−^ (Figure 2C). From the slope of the current-voltage relationship, Na^+^ conductance was estimated at G = 76.6 ± 4.1 pS, consistent with an open, cation-selective state.

**Figure 2.**
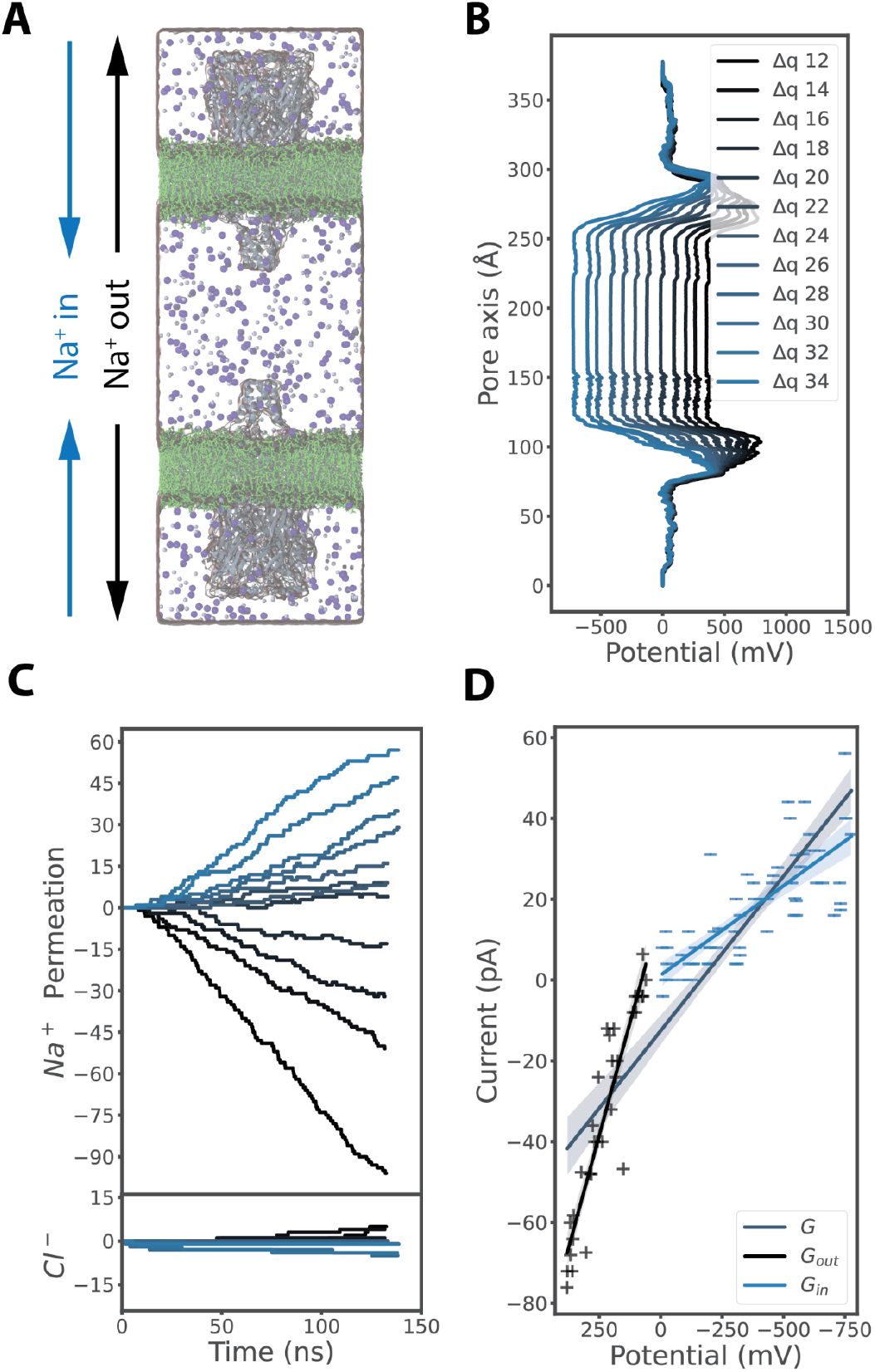
A conducting functional state under activating conditions. **A**. Snapshot of a computational electrophysiology simulation, with 150 mM NaCl (Na^+^, blue; Cl^−^, purple) surrounding two α7 nAChRs (gray) embedded in separate membrane patches (green) in an anti-parallel setup. **B**. Averaged electrostatic potential profiles along the membrane normal as shown in panel A, with various ion imbalances (differential of 12–34 charges, navy to light blue) between the two water compartments separated by the membranes. **C**. Permeation events over time for Na^+^ (above) or Cl^−^ ions (below) in individual computational electrophysiology simulations conducted at membrane potentials colored as in panel B (depolarized to polarized conditions, navy to light blue). Positive permeation events refer to ions passing from outer to inner compartments (“Na^+^ in,” panel A); negative values refer to ions passing the opposite direction. **D**. Current-voltage plot derived from computational electrophysiology simulations as shown in panel C. Solid lines show linear fits (with 95% confidence interval) to currents measured at depolarized potentials (navy +, G_out_ = 223.0 ± 16.5 pS), currents at polarized potentials (light blue –, G_in_ = 44.2 ± 4.4 pS), or the full dataset (steel, G_full_ = 76.6 ± 4.1 pS).

Interestingly, Na^+^ current-voltage relationships at polarized potentials fit a shallower conductance slope (G_in_ = 44.2 ± 4.4 pS) than those at depolarized potentials (G_out_ = 223.0 ± 16.5 pS) (Figure 2D). Indeed, the estimated depolarized (outward) Na^+^ conductance was remarkably similar to experimentally measured 192 pS (Noviello et al., 2021). This preference could reflect a modest outward rectification, or it could be a consequence of model or parameter bias, possibly underestimating inward flux. To test the robustness and determinants of this effect, we ran additional simulations of a single receptor in the presence of a hyperpolarized (–200 mV) or depolarized (+200 mV) external electric field. As expected, the structure determined under desensitizing conditions (PDB ID 7KOQ, with epibatidine alone) was effectively nonconductive in both conditions. Conversely, the structure under activating conditions exhibited comparable Na^+^ conductance as in our computational electrophysiology simulations, with lower inward than outward values (Figure 2S1).

Notably, deletion of the ECD produced a partial receptor with elevated conductance in both directions, and abolished the preference for outward flux, suggesting that this domain limits Na^+^ permeation particularly in the inward direction. In contrast, deletion of the ICD bundles resulted in conductance values comparable to wild-type in both directions, substantiating the importance of the ECD in suppressing Na^+^ efflux. Removing the sidechain at the tightest constriction in the ECD vestibule (E97A) also failed to relieve the apparent inhibition in either direction, indicating that conductance determinants are located elsewhere. A possible contributing factor was the presence of five Ca^2+^ ions, which were resolved at the ECD-TMD interface (Galzi et al., 1996; Noviello et al., 2021) and accordingly included in our computational electrophysiology experiments, as well as permeation calculations described below. Coordinated by acidic residues D41, D43, E44, and E172, these ions were distal to the conduction pathway, but raised the effective potential particularly in the outer half of the TMD pore (Figure 3S5). However, including these bound ions in our applied-field simulations only slightly elevated the conductance in both directions and did not substantially alter the preference (Figure 2S1), suggesting that local ion interactions in this region contribute little to the apparent ECD barrier to ion flow.

### State-dependent ion interactions from permeation free-energy profiles

To further characterize the functional states and ion interactions of each lipid-embedded cryo-EM structure, we next calculated permeation free-energy profiles for various ions and states, using the accelerated weight histogram (AWH) method (Lindahl et al., 2018, 2014). This approach to enhanced sampling is particularly suited to modelling the nonlinear conduction pathway of nAChRs, in which ions are expected to permeate the fenestrations between ICD helices (Figure 3S4B). The structure in the presence of inhibitory α-bungarotoxin (PDB ID 7KOO) featured a 40 kcal/mol free-energy barrier at the midpoint of the TMD. This apparent gate was centered at L247 (9′) (Figure 3A), a conserved residue known to form a hydrophobic gate in resting pLGICs. (Beckstein and Sansom, 2006; Ivanov et al., 2007). We henceforth considered this structure in the resting *state*. In contrast, under activating conditions (PDB ID 7KOX) the 9′ barrier decreased below 4 kcal/mol for Na^+^, K^+^, or Ca^2+^, comparable to a low barrier at the outer end of the ECD (Figure 3B, Figure 3S3A-B). Thus, based on its behavior in computational electrophysiology, applied-field, and ion-permeation simulations, this *activated*-state structure was apparently open.

**Figure 3.**
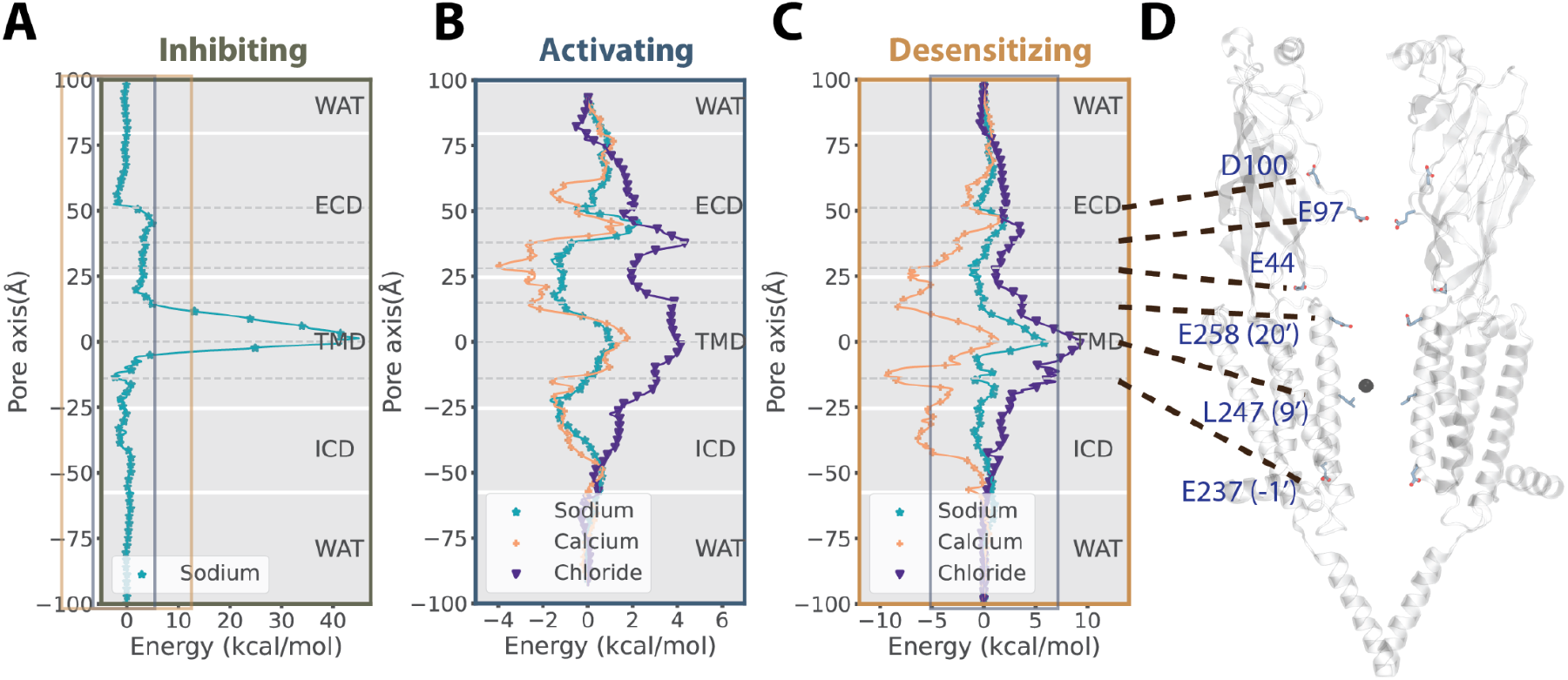
State-dependent ion interactions from permeation free-energy profiles. **A**. Free-energy profile of Na^+^ ion permeation (teal) through the lipid-embedded α7-nAChR structure under inhibiting conditions (PDB ID 7KOO). Dashed lines indicate key acidic residues, as well as the 9′ hydrophobic gate, facing the channel axis in the ECD and TMD. Solid boxes indicate zoom windows depicted in panels B (steel) and C (orange). **B**. Free energy profiles of Na^+^ (teal), Ca^2+^ (ochre), and Cl^−^ (indigo) permeation through the structure under activating conditions (PDB ID 7KOX), with key residues indicated as in panel A. All energy barriers are substantially reduced relative to inhibiting and desensitizing conditions, with the lowest barriers for Ca^2+^. **C**. Free energy profiles of ion permeation through the structure under desensitizing conditions (PDB ID 7KOQ), colored as in panel B, with key residues indicated as in panel A. Solid steel box indicates zoom window depicted in panel B. **D**. Model of the structure under activating conditions; for clarity, only two opposing subunits are shown. Key residues labeled in panels A–C are shown as sticks.

The α7 nAChR is known to be selective for cations (Séguéla et al., 1993). Indeed, Cl^−^ interactions were unfavorable throughout the activated-state conduction pathway, and particularly elevated relative to positive charges at 9′ and at the ring of E97 sidechains forming the tightest ECD constriction (Figure 3B). Conversely, favorable energy wells were observed for cations both at the outer end of the ECD near D100, and at the TMD-ICD interface (Figure 3B, Figure 3S3A). Notably, cations were directly coordinated by protein oxygen atoms at several positions in the ECD and at the outer and inner ends of the TMD pore (Figure 3S4A–B), possibly contributing to selectivity. As α7 nAChRs are particularly permeable to Ca^2+^ compared to other subtypes (Castro and Albuquerque, 1995), we sought to model Ca^2+^ interactions as accurately as possible, adopting recent parameters shown to be more accurate than default values in CHARMM36 (Zhang et al., 2020). Consistent with previous reports, we found this model to reasonably represent Ca^2+^ hydration (Figure 3S1) and to relieve potentially overestimated protein-Ca^2+^ interactions, particularly at the intracellular mouth of the pore (Figure 3S3D). In our calculations, aside from the peripheral binding site (near E44, Figure 3S2B), Ca^2+^ made several favorable interactions along the conduction pathway, including energy wells below -2 kcal/mol at the E97 constriction in the ECD, E258 (20′) in the outer TMD, and E237 (−1′) at the inner mouth of the TMD pore. Consistent with our free-energy profiles, experimental structures were reported with five Ca^2+^ ions peripheral to the ion permeation pathway (Figure 3S2B, Figure 3S4B), near the five symmetric E44 residues. To test the influence of bound Ca^2+^ on ion permeation, we also ran free energy calculations with Ca^2+^ at the five E44 sites in the structure determined under activating conditions (Figure 3S2). The presence of Ca^2+^ had no effect on the profile for Cl^−^ permeation. For Ca^2+^, it relieved the free energy well for further Ca^2+^ interactions at E44, but it did not substantially alter the permeation landscape elsewhere. Bound Ca^2+^ elevated the free-energy barrier for Na^+^ more broadly across the ECD-TMD interface, between E97 and E237, although the predominant barrier remained at the outer ECD around D100. Thus, inclusion of Ca^2+^ did not qualitatively alter the apparent permeation or selectivity of this structure, though local effects on ECD dynamics in the vicinity of E44 remain to be explored.

Under desensitizing conditions, the principal barrier to ion conduction was again found at the 9′ hydrophobic gate, and it is elevated relative to the activated state (Figure 3C, 3S3C). Notably, monovalent cations had to release at least one hydration water molecule in order to transit the 9′ gate in this structure, while in the activated state they retained a full hydration shell (Figure 3S4A, C, E). Ion-coordinating waters were further substituted by protein oxygen atoms at the inner mouth of the pore (E237, -1′; Figure 3S4E, F), though this position did not constitute a substantial free energy barrier (Figure 3C). For Ca^2+^, several free energy wells were especially pronounced under desensitizing conditions, with interaction energies below -7 kcal/mol at E44, E258 (20′), and E237 (−1′) relative to bulk solvent (Figure 3C). Accordingly, while Ca^2+^ ions were bound even more strongly in the pore of this structure than in the activated state, they also faced a more substantial (9 kcal/mol) barrier to transit the hydrophobic gate. Thus, along with the lack of conduction observed for this structure in applied-field simulations (Figure 2S1), we henceforth considered it a plausible *desensitized state*.

### State-dependent lipid interactions

With functional states assigned to each of the α7 nicotinic receptor structures, we further investigated lipid interactions in each state. Compared to the apparent resting and desensitized states, the activated-state experimental structure featured a distinctive translocation of the MX helix towards the membrane core (Figure 1A). This correlated to a compression of lipid density in the vicinity of the protein (Noviello et al., 2021). The timescales of lipid diffusion and remodeling in mixed membranes are typically prohibitive to all-atom simulations. Therefore, to test the extent and implications of this apparent membrane compression, we applied coarse-grained simulations to each structure modeled with Martini 2.2 (Marrink et al., 2007). To approximate the experimental system as closely as possible, each structure was embedded in a mixed bilayer with 20% cholesterol and simulated with protein backbone restraints for 20 µs (see Methods). We then analyzed protein-lipid interactions in the equilibrated systems during the final 5 µs of simulation time.

Membrane thickness proximal to the protein was strikingly state-dependent (Figure 4A). Whereas the resting and desensitized states were embedded in comparable local environments up to 42 Å thick, the activated state compressed the surrounding membrane to as little as 37 Å. After 20 µs coarse-grained simulation of the open structure, lipid heads from the inner leaflet translocated “up” towards the bilayer core (Figure 4C), increasing their interactions with the MX helix relative to the resting and desensitized states (Figure 4B). This effect dissipated within 60Å from the protein center (Figure 4S1), restoring the bulk membrane to roughly 40 Å thickness (Figure 4A, 4S1). The free-energy cost for this compression was estimated to be 0.7 or 0.4 kcal/mol for the resting-to-activated or desensitized-to-activated transitions, respectively (Methods).

**Figure 4.**
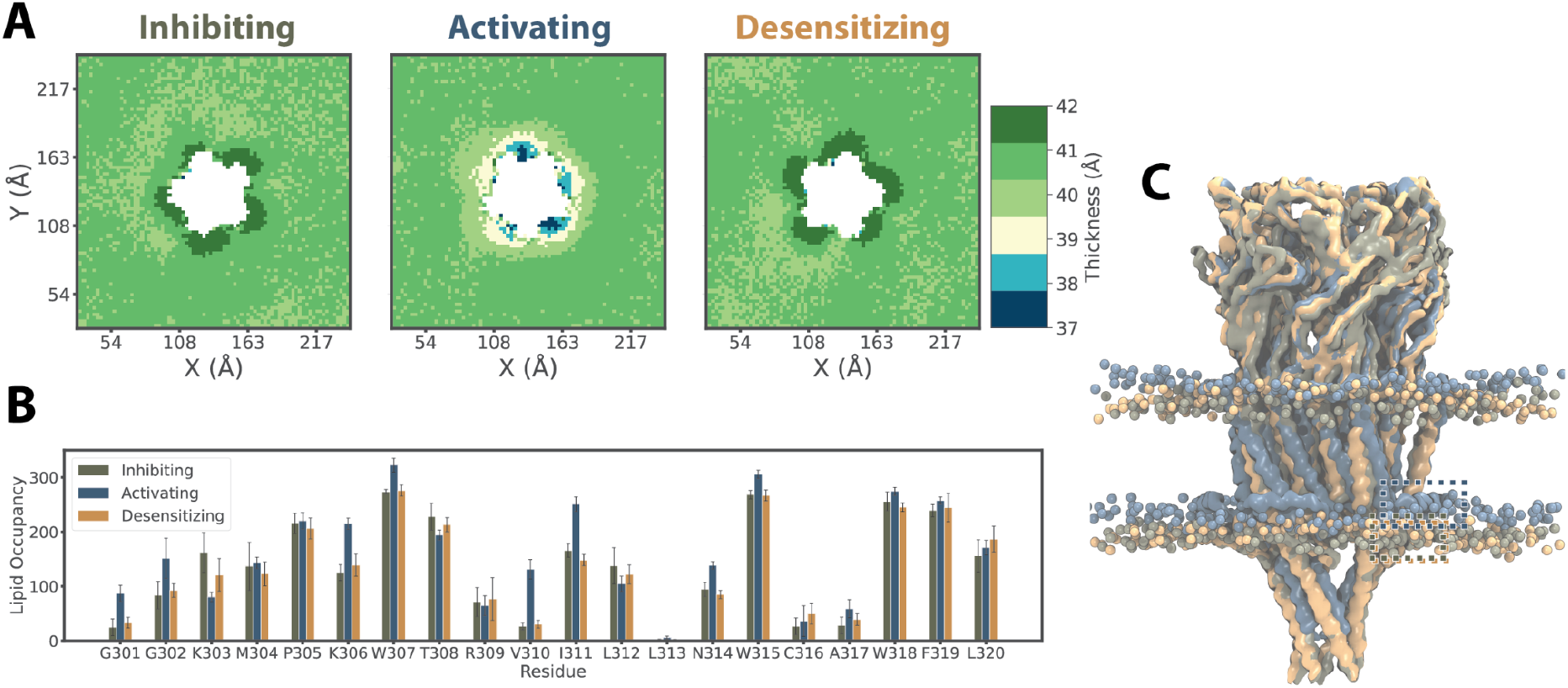
Local membrane compression in the activated state demonstrated by coarse-grained simulations. **A**. Membrane thickness, colored according to the scalebar at the far right, averaged over the last 5 μs from simulations of the lipid-embedded α7-nAChR structure under inhibiting (resting), activating (activated, center), or desensitizing (desensitized, right) conditions. The membrane thickness immediately proximal to the protein was comparable in the apparent resting and desensitized states (∼42 Å), while it was compressed in the activated state (37–39 Å). **B**. Occupancy of lipid interactions at each residue of the MX helix, showing increased contacts at several residues in the activated state (gray). **C**. Overlay of the last snapshots from simulations of the apparent resting (olive), activated (gray), and desensitized (orange) states, aligned on the ECD. Lipids of the inner leaflet are relatively displaced towards the membrane core by the distinct conformation of the activated state. Dashed boxes indicate positions of the membrane-peripheral MX helix in each structure. Only the membrane within 60 Å of the protein is shown for clarity.

Aside from the differences in bulk membrane properties described above, the coarse-grained simulations of α7-nAChR structures also indicated state-dependent interactions with specific lipids. Notably, cholesterol has been shown to modulate desensitization in nAChRs, although its role in the α7 subtype is unclear (Rankin et al., 1997). In simulations of the resting and activated states, cholesterol interacted with membrane-facing residues in the M1, M3, M4, and MX helices (Figure 5A–C), consistent with densities attributed to cholesterol in other recent α7 structures (Figure 5S2) (Zhao et al. 2021). In simulations of the desensitized state, cholesterol contacts shifted away from MX towards the outer-leaflet faces of M1 and M3. Moreover, cholesterol interacted extensively (>75% occupancy) with residue M253 (15′) in the pore-lining M2 helix in the desensitized state, via a cavity at the subunit interface (Figure 5A, D). This interaction was not observed in corresponding resting or activated-state structures, consistent with a role for intersubunit cholesterol binding in facilitating the distinctive desensitization profile of α7 nAChRs. Interestingly, this site overlapped with PNU density inside a partially desensitized/activated state of the α7 receptor in detergent (Zhao et al., 2021). These findings led us to further investigate possible PNU binding sites in the fully activated state.

**Figure 5.**
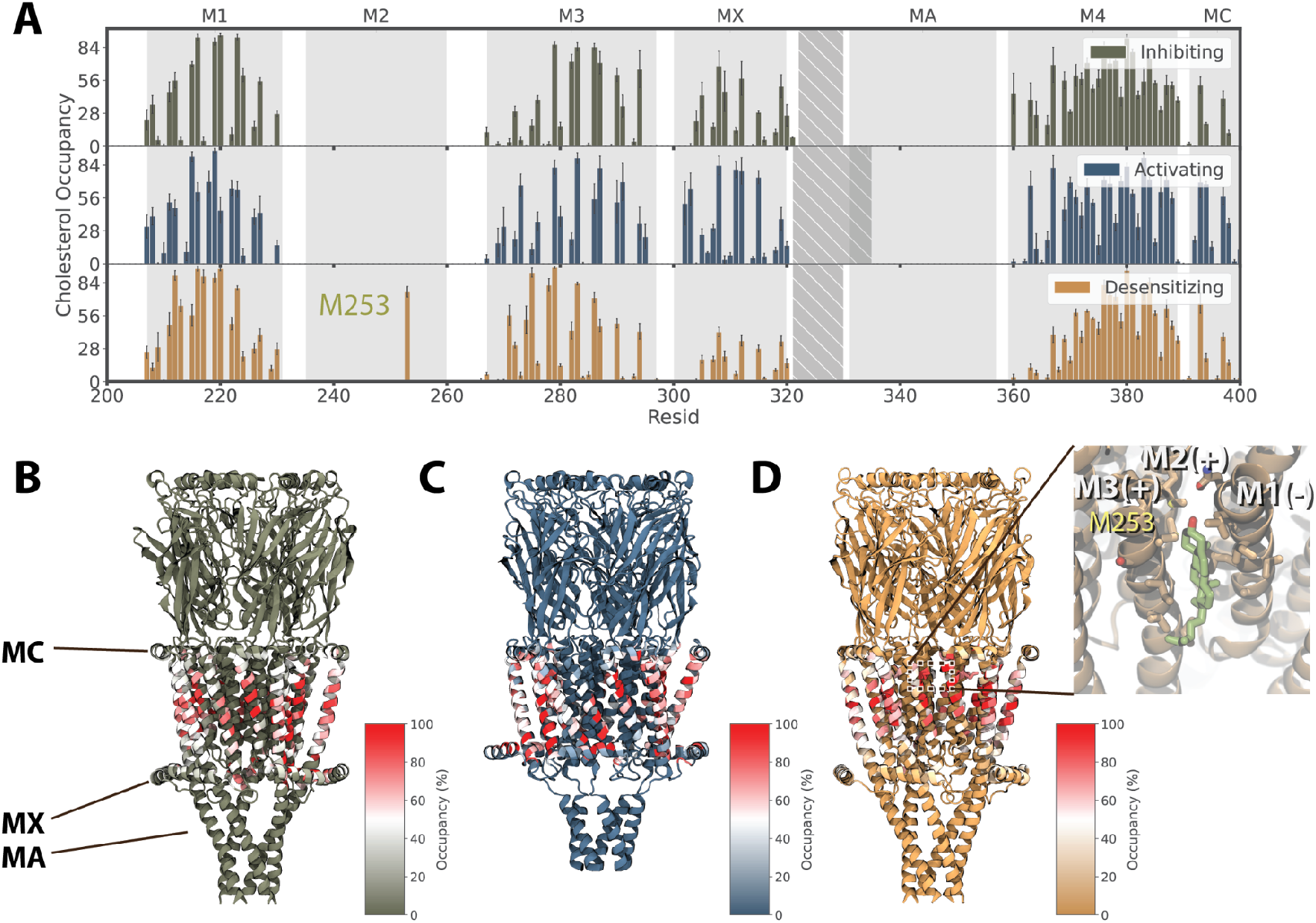
Altered cholesterol interactions in the desensitized state. **A**. Percent occupancy of cholesterol interactions with each residue of the α7 nAChR in 20-us coarse-grained simulations of structures determined under inhibiting (top), activating (middle), and desensitizing conditions (bottom). Whereas inhibiting and activating conditions were associated with comparable cholesterol interactions in the transmembrane core and MX helix, desensitizing conditions preferred interactions in the outer leaflet, including M2 residue M253. **B**. Cholesterol occupancies as in *A*, colored according to scalebar and mapped onto the experimental structure under inhibiting conditions, with key membrane-facing or peripheral helices labeled. **C**. Cholesterol occupancies as in *B* for the structure under activating conditions. **D**. Cholesterol occupancies as in *B* for the structure under desensitizing conditions. The inset shows cholesterol (green) and associated residues backmapped to atomic coordinates, including M253 from the principal M2 helix, in the upper-leaflet site preferred in this state.

### Allosteric potentiator PNU preferentially occupies an intersubunit site in the activated state

The presumed activated state was resolved in the presence of lipids with both the agonist epibatidine and positive allosteric modulator PNU (Noviello et al., 2021). However, despite saturating concentrations (200 μM), no PNU molecules could be definitively built in the cryo-EM density. To elucidate the binding mode of this modulator, we ran additional coarse-grained simulations of each experimental structure in the presence of PNU. Each system was built with ten PNU molecules placed randomly at a distance of 20 Å from the protein surface and simulated in quadruplicate for 20 μs using Martini 3 (Souza et al., 2021a, 2021b). This force field, recently reported to be better optimized for protein-ligand interactions (Souza et al., 2021b), recapitulated PNU properties optimized with quantum-mechanical or atomistic approaches beforehand (Figure 6S1).

Although average PNU densities were relatively diffuse and bound asymmetrically across the five subunits in simulations of the resting or desensitized states (Figure 6A), they consistently occupied sites in all five subunits facing the outer leaflet in the activated state symmetrically. Each of five primary, intersubunit outer-leaflet sites was bounded by the M2–M3 helices of the principal subunit, and M1–M2 of the complementary subunit (Figure 6B). Specifically, PNU bound to the activated-state structure preferentially assumed a pose 8 Å from M2 residue M253 and 5 Å from M3 residue M278 (center of mass distance), whereas interactions in other states sampled a wider range of poses. A secondary, peripheral outer-leaflet site was bounded by the M1, M3, and M4 helices of each individual subunit (Figure 6A, C). Although this site was visited with lower occupancy than the intersubunit site, PNU appeared to prefer an associated pose 5 Å from M1 residue S222 and 6 Å from M3 residue T273 in the activated state, in contrast to more distant and diffuse interactions in other states (Figure 6E). Mutations near both the intersubunit (M253L) and peripheral PNU sites (S222M, C459Y) were previously shown to disrupt PNU potentiation(Criado et al., 2011; daCosta et al., 2011; Young et al., 2008), indicating that both may contribute to binding of this modulator.

**Figure 6.**
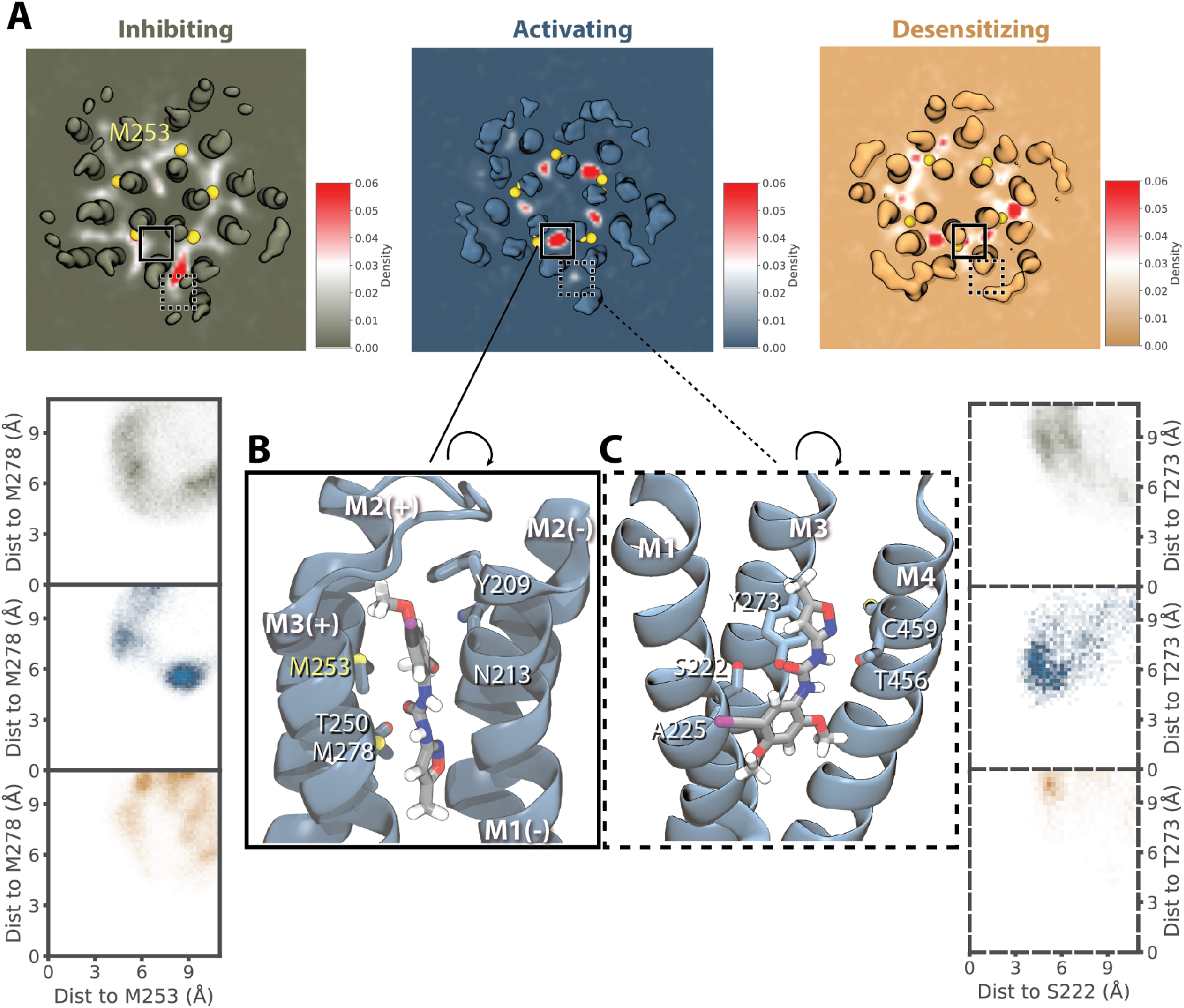
Allosteric potentiator PNU preferentially occupies an intersubunit site in the activated state. **A**. Average PNU density maps derived from quadruplicate 20 μs coarse-grained simulations, shown for a representative sliced through the transmembrane domain (Figure 6S3). A primary, intersubunit and secondary, peripheral site for PNU in the activated state are indicated by solid and dashed boxes, respectively. The intersubunit density of PNU was observed with relative symmetry over all five interfaces in the activated state (blue). **B**. PNU binding in the intersubunit site, backmapped to an all-atom model. The inset shows preferential occupation of a site bounded by residues M253 and M278 in the activated (middle) relative to resting (top) and desensitized states (bottom). **C**. Backmapped representation of PNU binding in the peripheral site, with inset as in *C* showing occupation of a pose bounded by S222 and T273.

Interestingly, recent structures of the α7 nAChR in detergent also showed PNU at an intersubunit outer-leaflet site (Zhao et al., 2021). Although the PNU site in this state, described as partially desensitized/activated, involved similar key residues as in our simulations, substantial differences in the helical backbone render them partially incompatible. If aligned on the M2 helices, the resolved pose for PNU—with its long axis parallel to the membrane plane—would clash with the outer M3 helix in our simulations (Figure 6S2), indicating that the primary pose described here is specific to the more expanded lipid-bound activated state.

To further test the role of the primary, intersubunit site in PNU binding, we ran additional coarse-grained PNU simulations of the activated structure with the mutation M253L. Despite the relatively conservative nature of this substitution, it consistently disrupted occupancy at the intersubunit site relative to wild-type (Figure 6S3). In M253L simulations, PNU instead occupied a novel inner-leaflet site, including contacts with M1 residue A225 (Figure 6S3). Interestingly, the substitution A225D was also previously shown to disrupt PNU potentiation (Young et al., 2008). In simulations of this A225D mutant, PNU again occupied the inner-leaflet site, making frequent interactions between its polar ureido group and the introduced aspartate at position 225 (Figure 6S3C). A double-mutant containing A225D and M253L exhibited a similar PNU as in both single mutants (Figure 6S3), consistent with a common structural effect of either disruption in the primary upper-leaflet site or enhanced binding in the lower-leaflet site.

Notably, our coarse-grained simulations indicated a pore-mediated pathway of PNU transit between intersubunit sites (Figure 6S4, Video 6S5). After entering an intersubunit site via the membrane, a PNU molecule could spontaneously transit the interface between M2 helices to enter the pore, then enter an equivalent site at another subunit interface. To partially quantify this phenomenon, we backmapped the activated-state coarse-grained system to atomistic resolution and used umbrella sampling to characterize this pore-mediated pathway. As expected, we found one free energy minimum within each intersubunit site, and a second in the open pore (Figure 6S4). A capacity for dynamic exchange between intersubunit sites could contribute to poor PNU resolution, even in the context of functionally relevant binding.

## Discussion

Understanding the gating cycle of ligand-gated ion channels requires comprehensive atomic details of functional endpoints, but also correlations with functional data and details of interactions with modulators and the lipid membrane components. With improvements in single-particle cryo-EM techniques, several pLGIC structures have now been resolved in the context of various ligands and lipidic environments. However, there are only a few cases in which a single construct has been reported in resting, activated, and desensitized states. In this context, the recent report of structures of the lipid-embedded α7 nAChR under inhibiting, activating, and desensitizing conditions provides an invaluable opportunity to mechanistic and dynamic modeling, not least due to the novel free energy profile and properties of the proposed activated state. A particularly critical challenge is the accurate assignment of functional states to experimental structures, for example by the application of molecular dynamics simulations of ion and lipid interactions, which we have pursued by combining electrostatic pore profiling, computational electrophysiology, applied electric fields, and enhanced sampling of ion permeation.

For the α7 nAChR, we found that simulations of the structure determined in presumed activating conditions—in the presence of both agonist (epibatidine) and potentiator (PNU)—produced single-channel conductance comparable to laboratory electrophysiology experiments (Noviello et al., 2021), which confirms its assignment as a plausible activated state. This assignment was further validated by simulations in the presence of an applied electrical field, and by the low barrier to cation permeation as determined by enhanced sampling. In contrast, the inhibited structure—determined with α-bungarotoxin—could be assigned to a resting state in simulations, with a major barrier to permeation at the central hydrophobic gate characteristic of these channels. A third structure, stably bound to an agonist (epibatidine) but impermeable to ions in our simulations, was accordingly assigned to a desensitized state. Interestingly, the predominant barrier to ion conduction in this structure remained at the central gate, albeit to a lesser extent than in the resting state. This free energy profile contrasted with those of desensitized gamma-aminobutyric acid-type A receptors, among others, in which the predominant gate was shown to shift to the intracellular end of the pore (Gielen et al., 2020). It remains to be determined whether this apparently distinct desensitization profile is shared by other nAChRs or in the larger pLGIC family.

Surprisingly, our simulations indicated a higher outward than inward conductance for Na+ ions in the activated state. In contrast, previous experimental work suggested a moderate inward rectification for α7 nAChRs (Alkondon et al., 1994; Forster and Bertrand, 1995). However, given that our simulations were performed in idealized computational conditions—lacking, for example, intracellular polyamines (Haghighi and Cooper, 1998) and substantial regions of the intracellular domain—it is not trivial to assess the biophysical or physiological relevance of this behavior. Simulations in the presence of an electric field indicated the apparent outward rectification could be attributed to differential inhibitory interactions in the ECD, without which conductance was enhanced to a similar level in both directions. This effect was not substantially modified by the presence or absence of the ICD, nor by an extracellular ring of glutamate residues (E97) at the tightest constriction in the channel pathway. An apparently structural set of modulatory Ca^2+^ ions, located peripheral to the conduction pathway in the ECD, also had little effect on conduction, though they remained stably bound throughout our simulations.

Both our computational electrophysiology and ion-permeation calculations demonstrated selectivity for cations over anions in the activated state. Notably, the free energy barrier for Ca^2+^ permeation was comparable to that for Na^+^. Accordingly, these channels should import substantial Ca^2+^ under physiological conditions, where intracellular buffering elevates the driving force relative to other cations. Using a recently refined Ca^2+^ model shown to approach the performance of some polarizable force fields (Zhang et al., 2020), we were able to identify local interaction sites for Ca^2+^ along the permeation pathway including acidic residues E97 and E258, previously shown to influence Ca^2+^ selectivity (Noviello et al., 2021). Moreover, with Ca^2+^ bound at its resolved sites in the ECD, interaction energies for Na+ were moderately elevated around the ECD-TMD interface. Meanwhile, Ca^2+^ interactions were more favorable relative to bulk solvent. Including the five structural Ca^2+^ ions in our simulation box approximated physiological extracellular concentrations of ∼2 mM, indicating that these preferences approximate a physiological profile. Abolishing Ca^2+^ binding at this site by the mutation D43A has been shown to decrease Ca^2+^ selectivity in electrophysiology experiments (Colón-Sáez and Yakel, 2014). It is consistent with a role for these bound ions in promoting Ca^2+^ permeation, possibly by a charge/space competition mechanism (Boda et al., 2002; Liu et al., 2021).

Coarse-grained simulations provided insights into the longer-timescale dynamics of lipid interactions and channel rearrangements associated with gating. Most dramatically, the activated state was associated with local compression of the membrane relative to resting or desensitized states, and increased interactions of the intracellular MX helices with membrane lipids. It is interesting to consider whether MX-lipid interactions may contribute to alleviating the free energy penalty of compressing the lipid bilayer. A similar relative motion of the MX helices towards the TMD has been reported in 5-HT3 receptors (Basak et al., 2019; Polovinkin et al., 2018), indicating such a conformational shift could be linked to a conserved mechanism of activation. Conversely, the desensitized state was associated with differential cholesterol interactions compared to resting or activated states, providing testable hypotheses for future work to refine our understanding of the established cholesterol dependence of gating in these channels (Barrantes, 2012, 2004; John E.Baenziger, Jaimee A.Domville, J.P. Daniel Therien, 2017). Moreover, we were able to apply the improved small-molecule interaction properties of the Martini 3 force field (Souza et al., 2021a, 2021b) to identify the activated-state binding pose of PNU, which was present but not resolvable in the lipid-nanodisc cryo-EM structure. In our simulations, PNU spontaneously transitioned between equivalent binding sites at the five subunit interfaces, suggesting dynamic behavior that could contribute to relatively poor resolution of the average experimental electron density. Interestingly, PNU occupied a similar region, but in a substantially different orientation, in a proposed intermediate state of the α7 receptor reported in detergent after initiation of this simulation work. Coarse-grained simulations may prove similarly valuable in characterizing state-dependent interactions particularly of lipophilic modulators as alternative structural models become increasingly available.

Taken together, the atomic-resolution and coarse-grained simulations in this work support a structurally detailed mechanism for a basic three-state gating cycle in receptors (Figure 7). Binding of the agonist in the ECD promotes opening of a cation-, and specifically Ca^2+^-, permeable pore, accompanied by local compression of the bilayer and increased contacts of the MX helices with the lower membrane leaflet. With continued agonist exposure, the channel would be expected to transition rapidly to a desensitized state, with a partially contracted pore and a shift in preferential cholesterol binding from a lower-to middle-leaflet site. However, binding of PNU— possibly in competition for this middle-leaflet site—relatively stabilizes the activated state, opposing desensitization. Although further structure-function work may refine this mechanism, including the likely contribution of additional resting, activated, desensitized, and intermediate states, this mechanism is consistent with the broad functional properties of the α7 receptors, and illustrates the evolving utility of molecular dynamics simulations in annotating and interpreting structural data for pharmacologically important systems.

**Figure 7.**
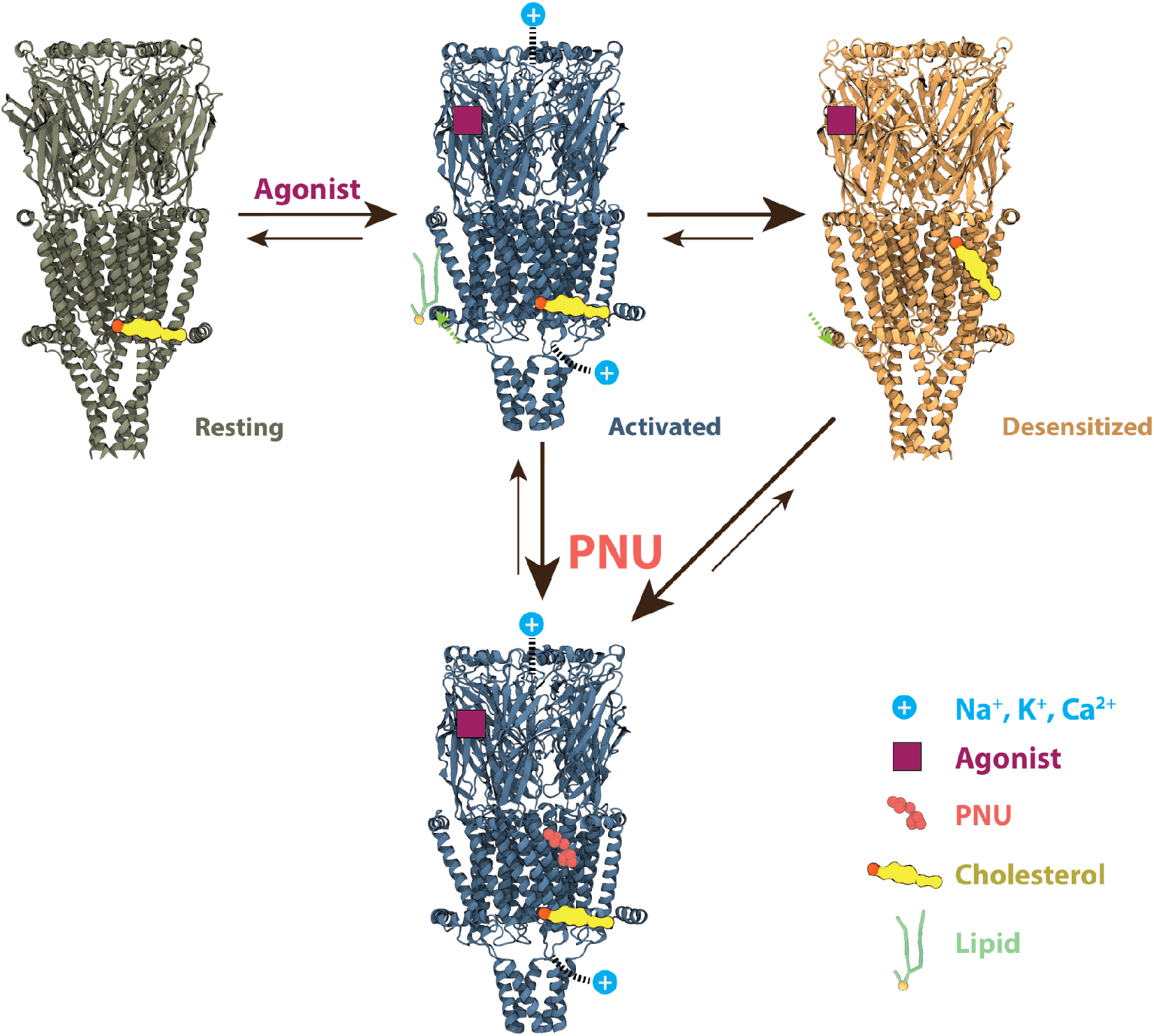
Proposed mechanism for α7 gating in the presence of agonist, with or without PNU. Starting from a resting state (olive), binding of agonist (purple) promotes transition to a Ca^2+^-permeable activated state (blue, top) with local membrane compression and increased contacts of the MX helix with lipids (green). This state can be relatively stabilized by binding of PNU (salmon) to a site near the middle of the TMD (blue, below). In the absence of PNU, the activated state transitions rapidly to a desensitized state (orange) with cholesterol (yellow) relocated to the middle-TMD site.

## Methods

### General all-atom MD-simulation setup

Coordinates for α7-receptor cryo-EM structures determined under apparent inhibiting, activating, and desensitizing conditions (PDB IDs 7KOO, 7KOX, 7KOQ respectively) were used as starting models for all simulations. Where present, the α-bungarotoxin, epibatidine, and Ca^2+^ ions were placed as in the deposited structures. The CHARMM36M (Huang et al., 2017) (July 2020 version) force field was used to describe each protein, which was embedded in a bilayer of 400 1-palmitoyl-2-oleoyl-sn-glycero-3-phosphocholine (POPC) molecules and solvated in a cubic box using CHARMM-GUI (Jo et al., 2008; Lee et al., 2016). The TIP3P (Jorgensen et al., 1983) water model and NaCl were added to bring the system to neutral charge and an ionic strength of 0.15 M. In total, 212 chloride ions, 242 sodium ions, and 76,871 water molecules were added into a 12.8 nm * 12.8 nm * 18.6 nm simulation box.

Atomistic simulations were performed with either default Ca^2+^ parameters or recently revised versions (Zhang et al., 2020) as indicated. Small-molecule parameters for epibatidine and PNU were generated with CGenFF (Vanommeslaeghe et al., 2010) and optimized with FFParam (Kumar et al., 2020) together with Psi4 (Smith et al., 2020) as the quantum chemistry backend. The modified version of the forcefield can be found as indicated under Data Availability.

All simulations were performed with GROMACS 2020 (Páll et al., 2020). Default settings of the CHARMM36 force field were applied during energy minimization and equilibration. Each system was energy-minimized, then equilibrated for 250 ps with a constant number of particles, volume, and temperature when both protein and lipid molecules were restrained. Each system was then equilibrated with a constant number of particles, pressure, and temperature for 40 ns during which the position restraints on the protein were gradually released. Weak thermostats and barostats (Berendsen et al., 1984) were used to model the system at 300K and 1 bar during relaxation, and bond lengths were constrained using the LINCS (Hess et al., 1997) algorithm.

### Computational electrophysiology

The equilibrated simulation box was used as the starting configuration. A second box was then translated, rotated 180 degrees, and merged with the first to generate an antiparallel alignment setup with comparable water content in the two membrane-delineated compartments. A small offset (0.08 nm) was included to ensure no collapse at the edges of the two simulation boxes. 20 ns of extra equilibration were included after energy minimization of the new double-bilayer simulation box. The ion permeation pathway was defined by two 1.2 nm-radius cylinders centered at residue 247 (defined as the compartment boundary) extending 7.5 nm towards the channel ECD and 5 nm towards the channel ICD. The swapping frequency was set to 100, the threshold to 1, and the coupl-step to 10. A comprehensive system setup script and the corresponding mdp file are available as indicated in Data Availability.

Potential differences were generated by varying sodium ion concentrations in the aqueous compartments, keeping the chloride ion concentration constant. Each simulation was run for 140 ns with protein Cα atoms restrained to prevent deviations from the starting state. The potential was quantified with the GROMACS potential tool in double precision. Net-zero charge of groups was assumed to improve accuracy. During calculations, a translation along the membrane normal z by half of the box length was applied to eliminate potential inaccuracies. To calculate single-channel conductance, ionic current as a function of the potential difference was determined within 20-ns time windows, with 10-ns overlap between consecutive windows. For electric-field simulations, a potential of ±200 mV was applied to the single-bilayer system. Modified channels without ECD, without ICD, or with the mutation E97A were embedded into the bilayer and equilibrated as previously described. The conductance was quantified by the corresponding current, i.e., the number of ions permeating during the simulations, divided by the applied potential.

### Accelerated weight histogram

AWH methods have been widely applied to study the ion-permeation free energy profiles in other channels (Kim et al., 2020; Lindahl et al., 2018, 2014). Unlike umbrella sampling, AWH does not have defined initial configurations, but flattens free energy barriers along the reaction coordinate to converge to a freely diffusing ion. This method was used to calculate free energy profiles along the pore axis for Na^+^, K^+^, Ca^2+^, and Cl^−^. For each equilibrated cryo-EM structure, one ion was additionally placed in the center of the pore around E258; a flat-bottomed restraint was applied to the ion to keep a radial distance below 20 Å from the pore axis. An independent AWH bias with a force constant of 12,800 kJ/mol/nm^2^ was applied to the center-of-mass z-distance between the selected ion and residue 247, with a sample interval across more than 95% of the box length along the z axis to reach periodicity. Semi-isotropic pressure coupling was used to keep the pressure to 1 bar where the compressibility along the z axis was set to 0 to ensure a constant sampling coordinate. A total of 16 walkers sharing bias data and contributing to the same target distribution were simulated for >100 ns until the PMF profile converged in more than 10 ns.

### Coarse-grained simulations

Coordinates for the same three α7-receptor cryo-EM structures (PDB IDs 7KOO, 7KOX, 7KOQ) without ions, ligands, or glycans were coarse-grained, through the representation of roughly four heavy atoms as a single bead, using Martini Bilayer Maker (Hsu et al., 2017) in CHARMM-GUI (Jo et al., 2008). The protein was embedded in a symmetric membrane containing 20% cholesterol (CHOL), 16% POPC, 24% PIPC, 4% POPE, 12% PIPE, 4% POP2, 12% PIPI, 4% POPA, and 4% PIPA, which approximates the soy-lipid mixture used for experimental reconstitution of this receptor (Noviello et al., 2021) (PC: phosphatidylcholine, PE: phosphatidylethanolamine, P2: phosphatidylinositol bisphosphate, PI: phosphatidylinositol, PA: phosphatidic acid; PO corresponds to a C16:0/18:1 lipid tail, while PI corresponds to a C16:0/18:2 lipid tail). In total, 2,500 lipids were inserted in a 27 nm * 27 nm * 20 nm simulation box, constituting ∼130,000 total beads including water and ions. After energy minimization and equilibration in CHARMM-GUI, simulations were run with the protein restrained for 20 μs in GROMACS 2020 to allow lipid convergence, using Martini 2.2 and 2.0 parameters for amino acids and lipids (Marrink et al., 2007), respectively.

The penalty of membrane compression was quantified (Ursell et al., 2007) as

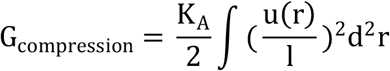

where *K*_*A*_ is the bilayer area stretch modulus (∼60 kT/nm^2), *u(r)* is the deformation from the unperturbed leaflet thickness, and *l* is the unperturbed leaflet thickness.

By summing the grid-based average membrane-thickness penalty,

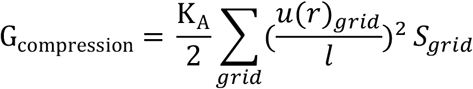

the free energy difference from membrane compression penalty can be calculated between three functional states:

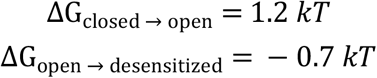

### Simulations with PNU

The coarse-grained model of PNU was built using the CG builder tool (https://jbarnoud.github.io/cgbuilder/). CG bead types and bonded parameters were assigned according to the Martini 3 forcefield (Souza et al., 2021a). Parameters were then optimized with Swarm-CG (Empereur-Mot et al., 2020) to fit the all-atom parameters. Permeation PMFs along the POPC membrane bilayer were then profiled with umbrella sampling (described below) for validation. After converting the entire system to Martini 3, ten PNU molecules were placed randomly into the simulation box (1 nm away from the protein). The backbone beads of the protein were restrained for better sampling. Four replicates of each system were then run for 20 μs each.

### Umbrella sampling

An output frame from coarse-grained simulations, in which one PNU was bound to the intersubunit binding site, was used as an initial configuration and backmapped to atomistic coordinates. The center-of-mass x/y-plane distance of PNU from the five position-253 residues, which would be in the middle of the pore, was used as the pulling coordinate. The “distance” was set to be the pulling-coordinate geometry. PNU was pulled either into the pore or out to the membrane at a rate of 0.0005 ns/ps with a force constant of 10000 kJ/mol/nm^2^ to generate initial configurations, with an interval of 0.04 nm spanning 0 nm to 3.7 nm. In total, 97 umbrella-sampling windows were simulated for 100 ns. The PNU was position-restrained with a flat-bottomed potential to keep it in a 2-nm layer parallel to the x/y plane for convergence. The weighted histogram analysis method (WHAM) (Kumar et al., 1992) was used to analyze the results. To measure permeation of PNU across the POPC bilayer, a similar setup was applied to either the all-atom system in CHARMM36M or the coarse-grained system in Martini 3. Umbrella-sampling windows with an interval of 0.05 nm were generated where the center-of-mass z distance between the PNU and the membrane was pulled with a “direction” geometry. The PBC atom of the membrane was set to -1 to turn on cosine weighting.

### Visualization and analysis tools

Visualizations were created in VMD (Humphrey et al., 1996); most analyses were performed with GROMACS and MDAnalysis (Michaud-Agrawal et al., 2011) and plotted with RainCloudPlot (Allen et al., 2019) and matplotlib (Hunter, 2007). For pore-radius calculation and visualization, CHAP (Rao et al., 2019) was used. G_elpot (Kostritskii et al., 2021) was used for quantifying the electrostatic potential along the channel with or without Ca^2+^. For the coarse-grained simulations, MemSurfer (Bhatia et al., 2019) was used to quantify membrane thickness and PyLipid (Song et al., 2021) was used to measure and map the occupancy and residence time of different lipids onto the protein.

## Acknowledgements

This work was supported by grants from the Knut and Alice Wallenberg Foundation, the Swedish Research Council (2021-02433, 2021-05806), the Swedish e-Science Research Centre, the BioExcel Center of Excellence (EU 823830), and the US National Institutes of Health (NS120496). Computational resources were provided by the Swedish National Infrastructure for Computing (SNIC 2021/3-39, 2021/37-14) and PRACE project 2020225362 at CSCS, Switzerland.

## Data Availability

Modified forcefield parameters, system-setup scripts, and simulation parameter files are available on Zenodo: 10.5281/zenodo.5782906.

## Supporting Information

**Figure 2S1.**
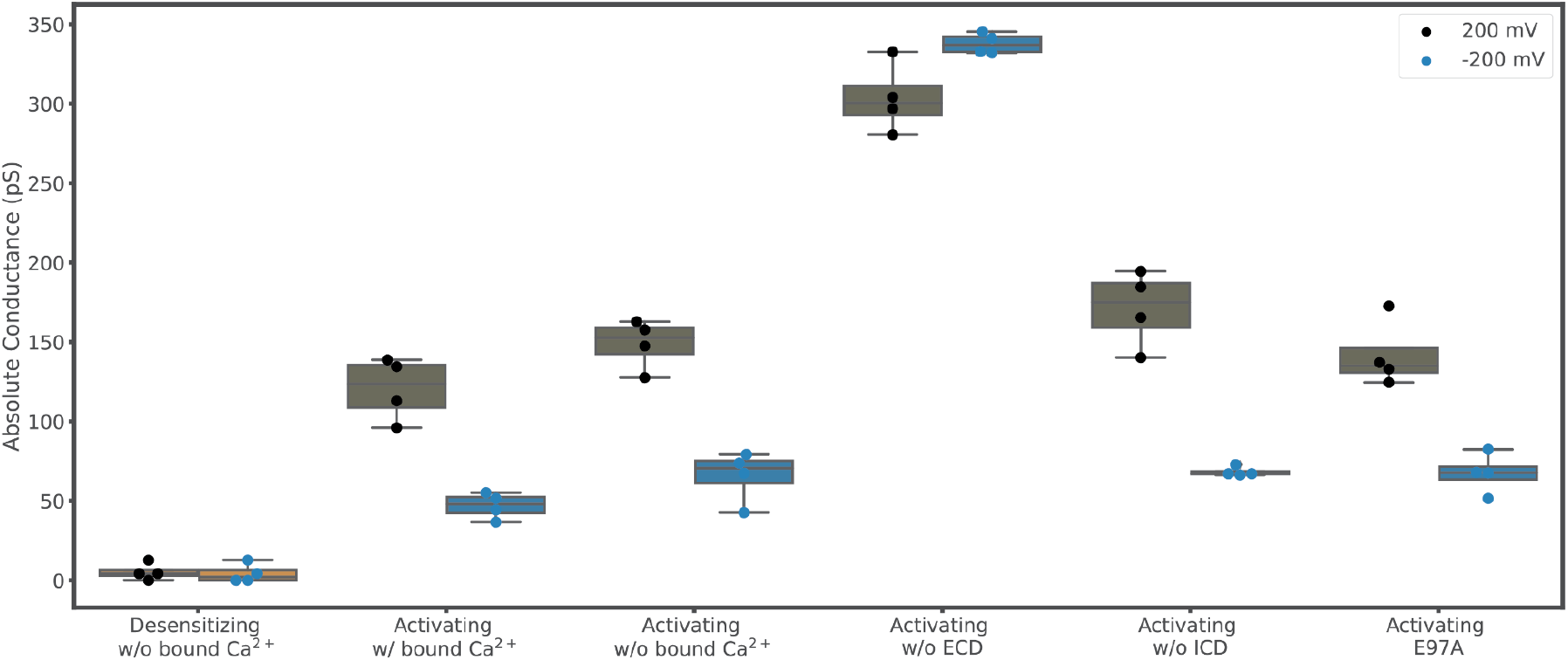
Applied electric field simulations support a conducting functional state in activating conditions. Average conductance measured at 200 mV (gray) and -200 mV (blue) potentials for structures determined under (left–right) desensitizing conditions, activating conditions (with or without bound Ca^2+^), activating conditions without the extracellular domain, activating conditions without the intracellular domain, or activating conditions with mutation E97A.

**Figure 3S1.**
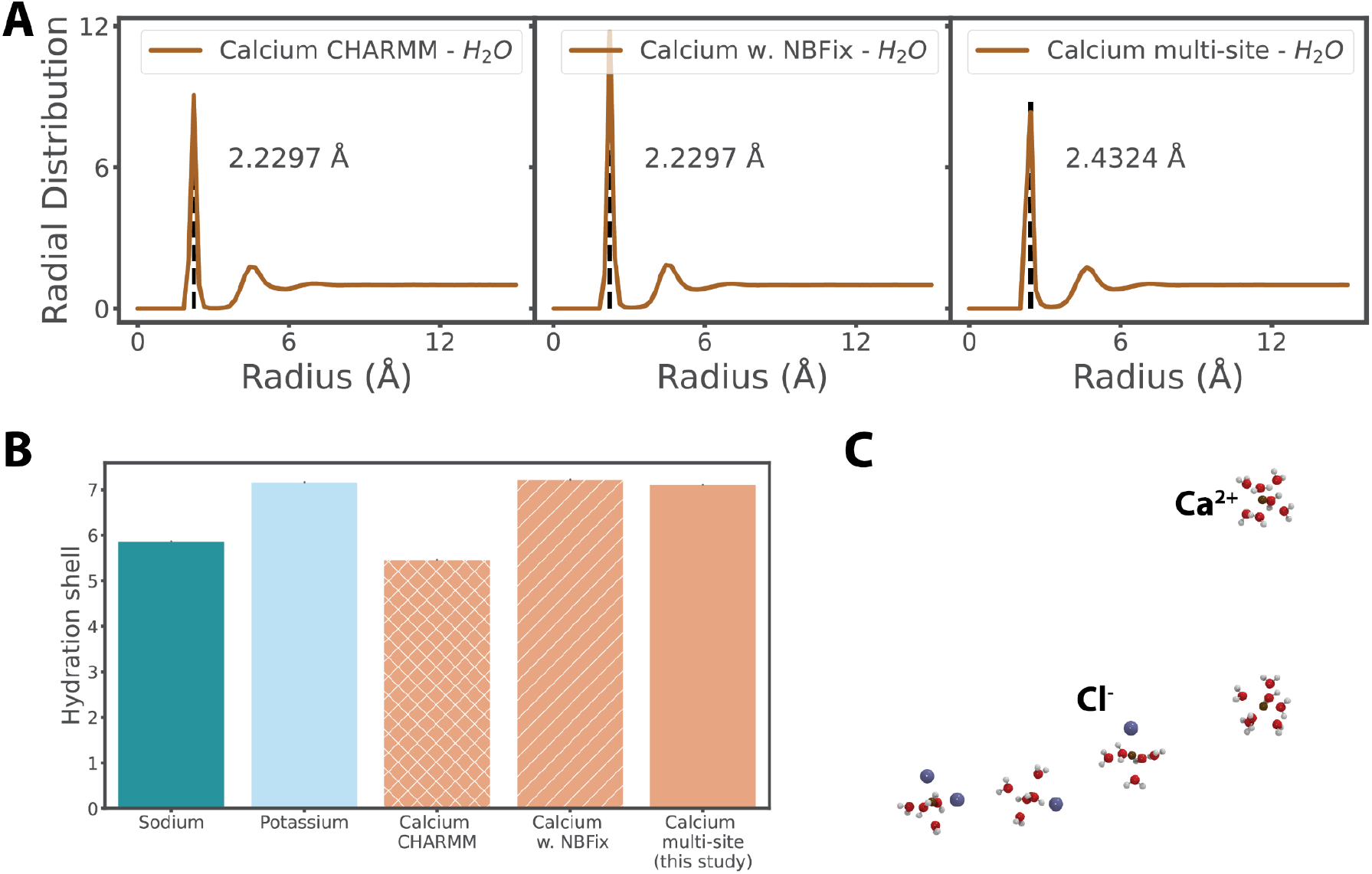
Comparative performance of Ca^2+^ models. **A**. Radial distribution function of ion-water oxygen atom pairs using the (left–right) CHARMM36, revised CHARMM36 with NBFix, and multi-site (CAM) Ca^2+^ models. **B**. Hydration shells of various ion species. **C**. Snapshots of solvation shells involving water (red) and Cl^−^ ions (indigo) surrounding Ca^2+^ ions (blue) using the original CHARMM36 Ca^2+^ model.

**Figure 3S2.**
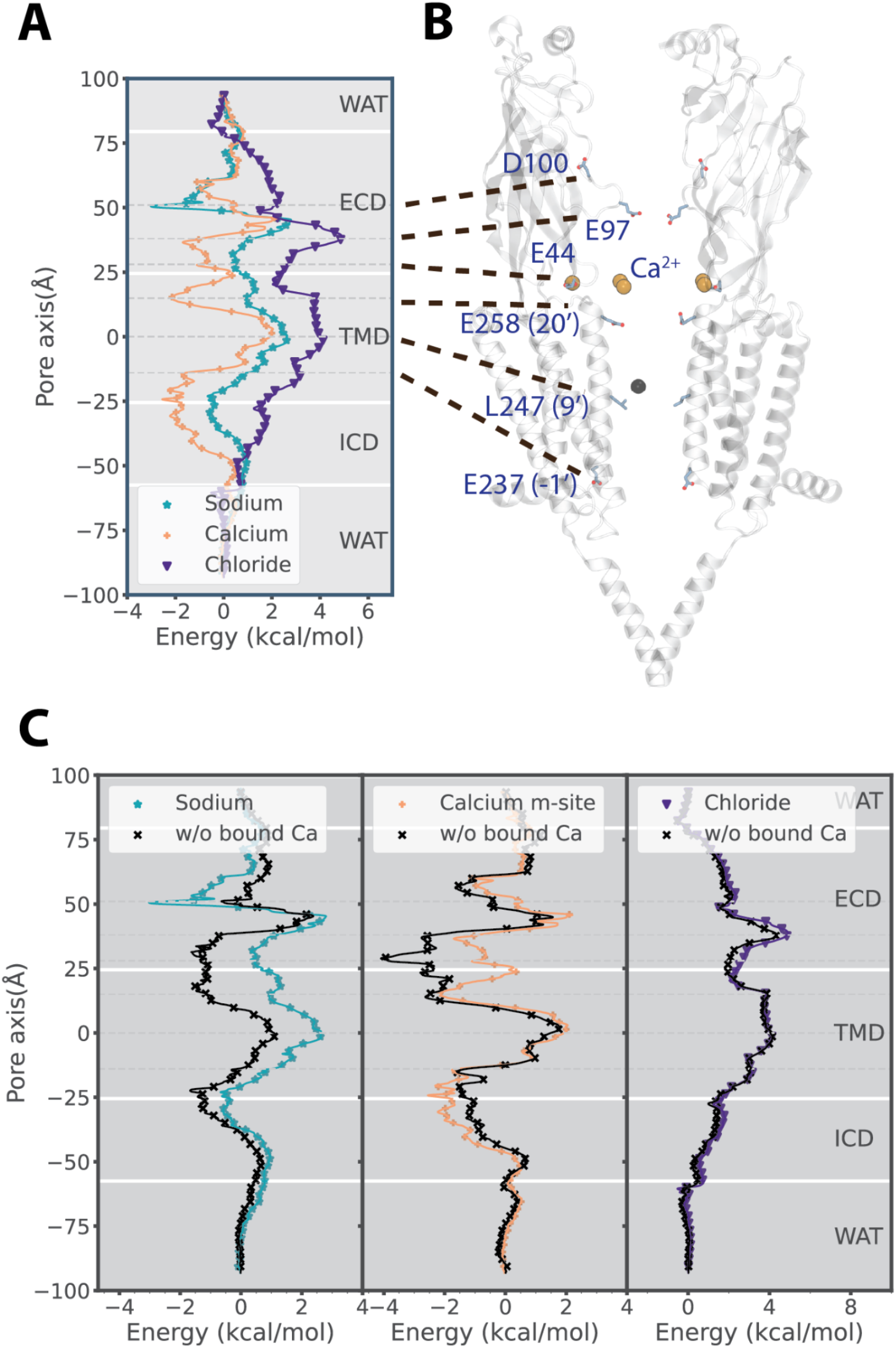
Bound Ca^2+^ perturbs permeation free-energy profiles in the activated state. **A**. Free-energy profiles of Na^+^ (teal), Ca^2+^ (ochre), and Cl^−^ (indigo) permeation through the structure determined under activating conditions (PDB ID 7KOX) with resolved Ca^2+^ ions explicitly included in simulations. **B**. Protein model of the structure determined under activating conditions; for clarity, only two opposing subunits are shown. Key residues labeled in panel A are shown as sticks, along with five Ca^2+^ ions (ochre) resolved in the ECD. **C**. Comparison of free-energy profiles with or without bound Ca^2+^ for the structure determined under activating conditions.

**Figure 3S3.**
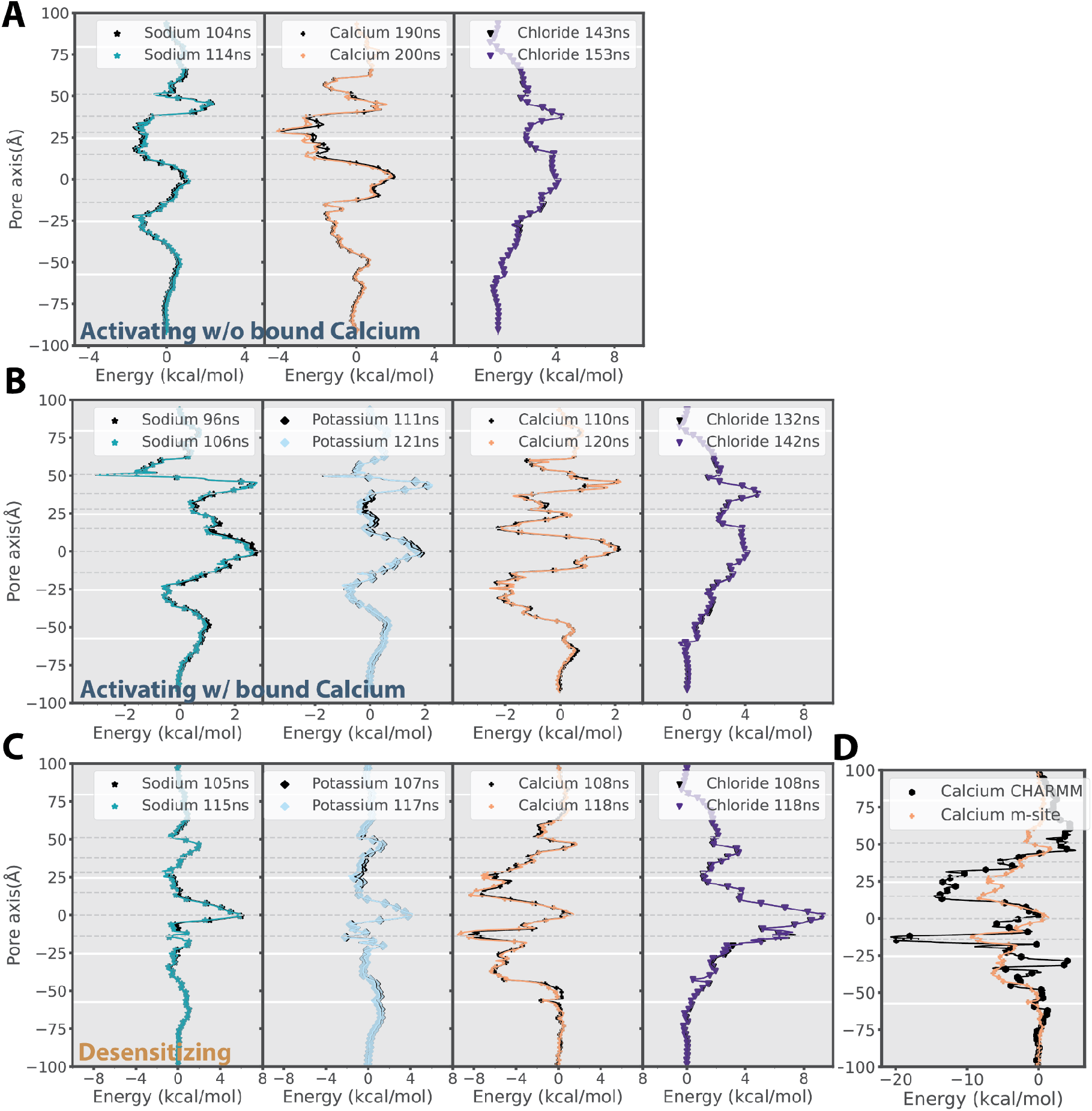
Convergence of permeation free-energy profiles. **A**. The free energy profiles of ion permeation for the activated state without bound Ca^2+^. **B**. The free energy profiles of ion permeation for the activated state with bound Ca^2+^. **C**. The free energy profiles of ion permeation for the desensitized state. **D**. Comparing the free energy profiles of Ca^2+^ permeation for the desensitized state with the CHARMM36 parameter and the revised multi-site Ca^2+^ model (CAM).

**Figure 3S4.**
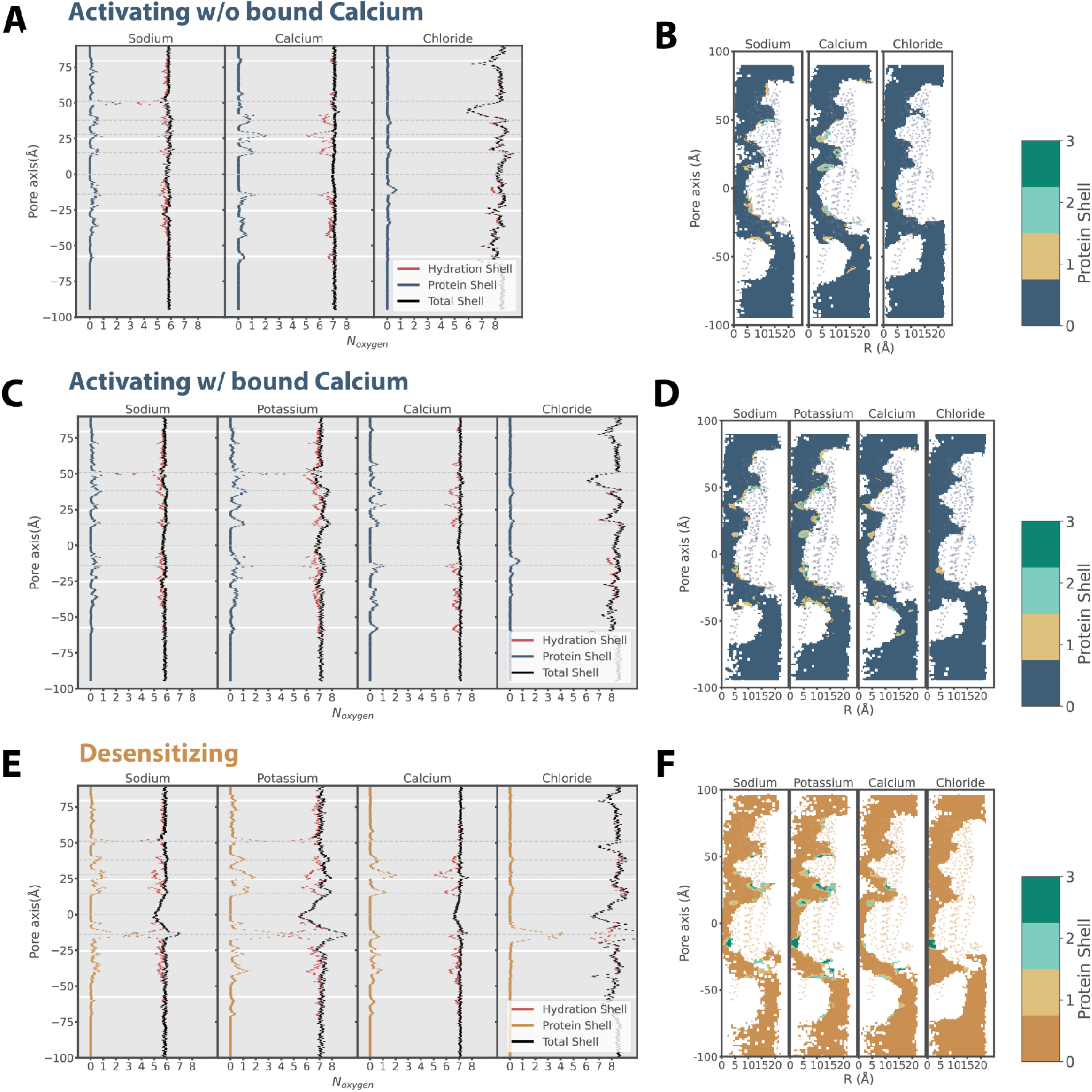
Water/protein coordination for various ions. **A**. The average number of coordinated oxygen atoms from water/protein as a function of position along the pore axis in the activated state without bound Ca^2+^. **B**. The heat map of the average number of coordinated oxygen atoms from protein in the activated state without bound Ca^2+^. **C**. The average number of coordinated oxygen atoms from water/protein as a function of position along the pore axis in the activated state with bound Ca^2+^. **D**. The heat map of the average number of coordinated oxygen atoms from protein in the activated state with bound Ca^2+^. **E**. The number of coordinated oxygen atoms from water/protein as a function of position along the pore axis in the desensitized state. **F**. The heat map of the average number of coordinated oxygen atoms from protein in the desensitized state.

**Figure 3S5.**
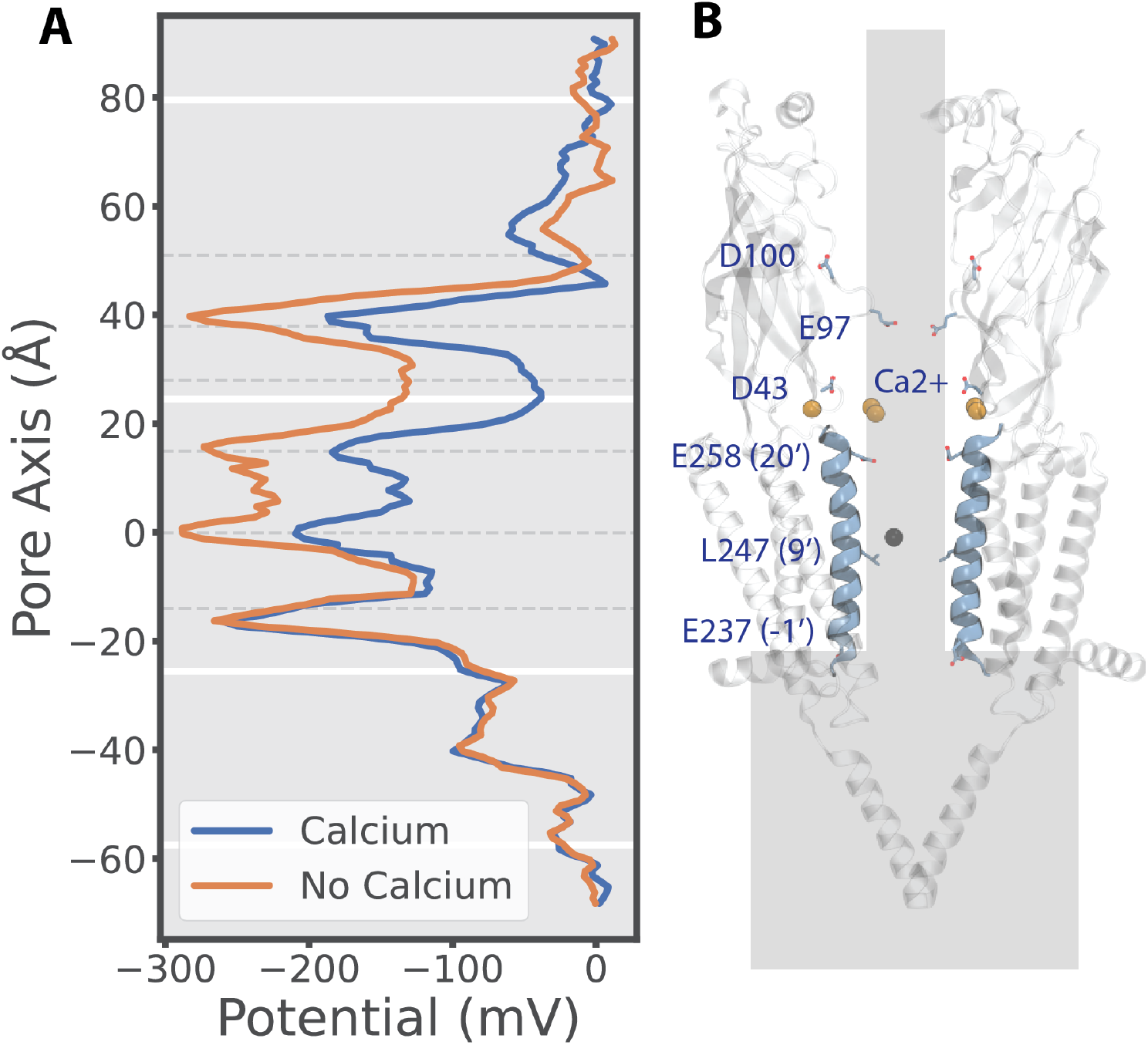
Electrostatics along the channel pore. **A**. The electrostatics along the channel with or without Ca^2+^ calculated with g_elpot (Kostritskii et al., 2021). **B**. The electrostatics calculated region mapped onto the structure under activating conditions. Key residues are shown as sticks, along with five Ca^2+^ ions (ochre) resolved in the ECD.

**Figure 4S1.**
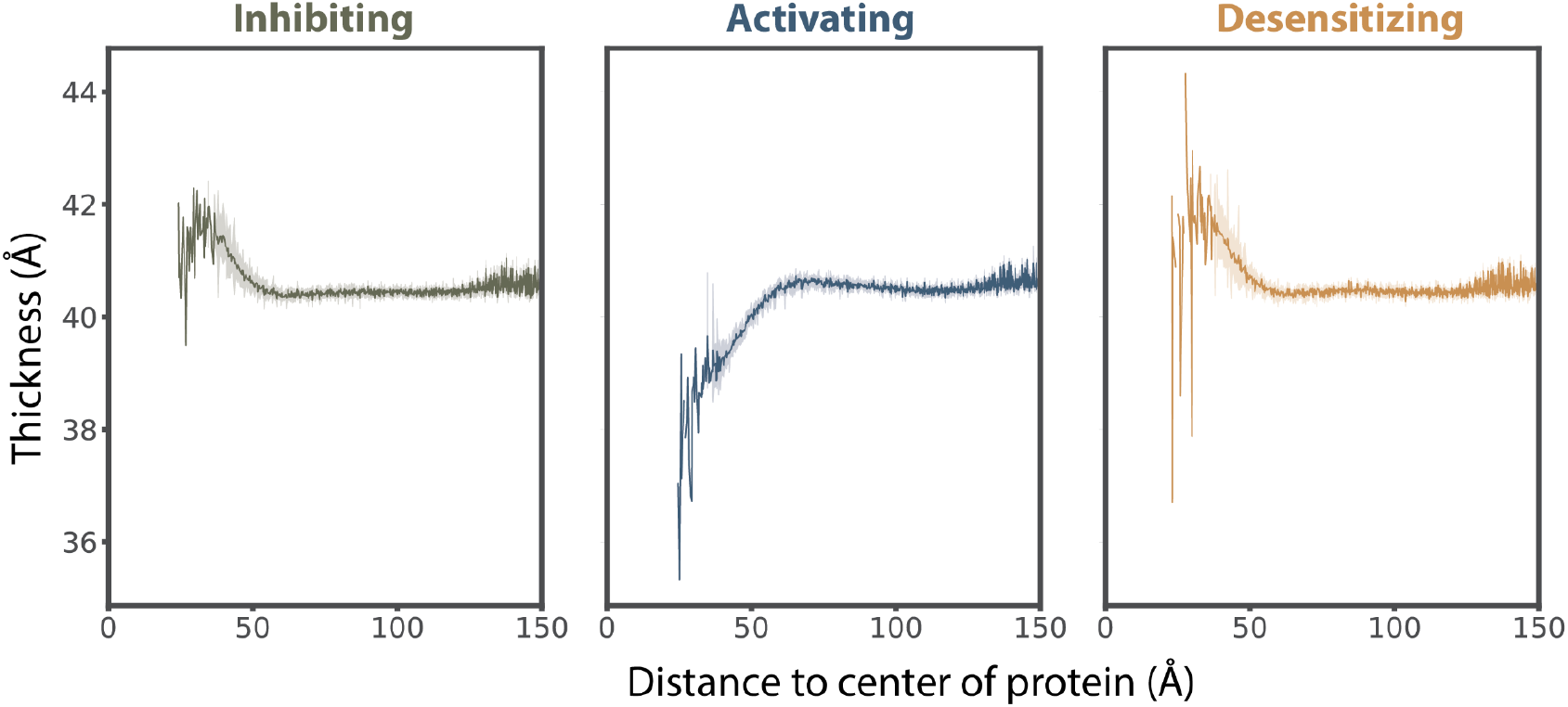
Quantification of membrane thickness. The perturbed membrane thickness differences averaged over the last 5 μs simulations dissipated within 60 Å from the protein for the lipid-embedded α7-nAChR structure under inhibiting (resting, left), activating (activated, center), or desensitizing (desensitized, right) conditions.

**Figure 5S1.**
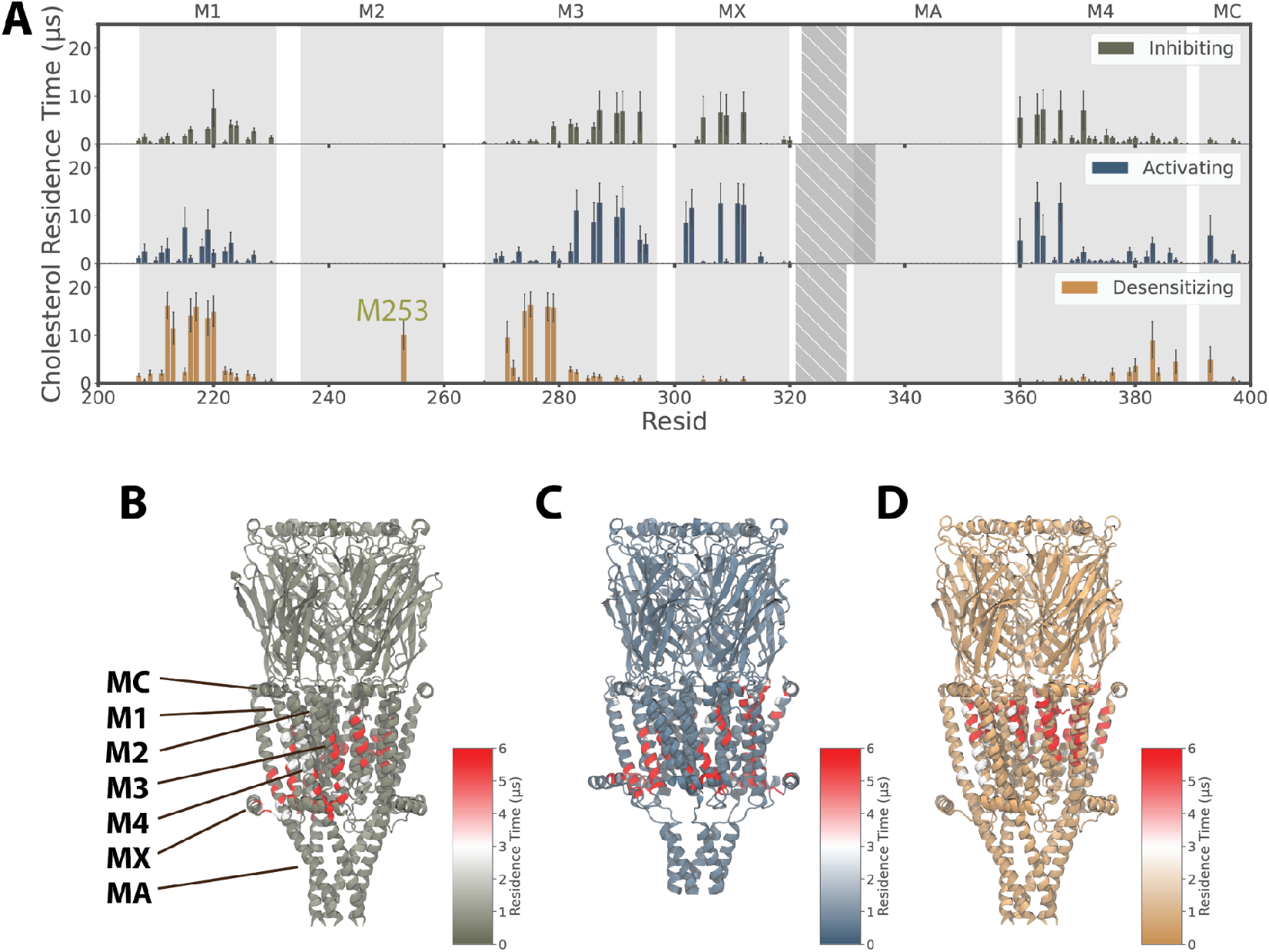
Residence time of cholesterol interactions in the desensitized state. **A**. The residence time of cholesterol interactions with each residue of the α7 nAChR in 20-us coarse-grained simulations of structures determined under inhibiting (top), activating (middle), and desensitizing conditions (bottom). While the resting and activated states shared a similar cholesterol interaction site in the lower part of the transmembrane domain the desensitized state shifted the cholesterol interactions upwards. **B**. Cholesterol residence time as in *A*, colored according to scalebar and mapped onto the experimental structure under inhibiting conditions, with key membrane-facing or peripheral helices labeled. **C**. Cholesterol residence time as in *B* for the structure under activating conditions. **D**. Cholesterol residence time as in *B* for the structure under desensitizing conditions.

**Figure 5S2.**
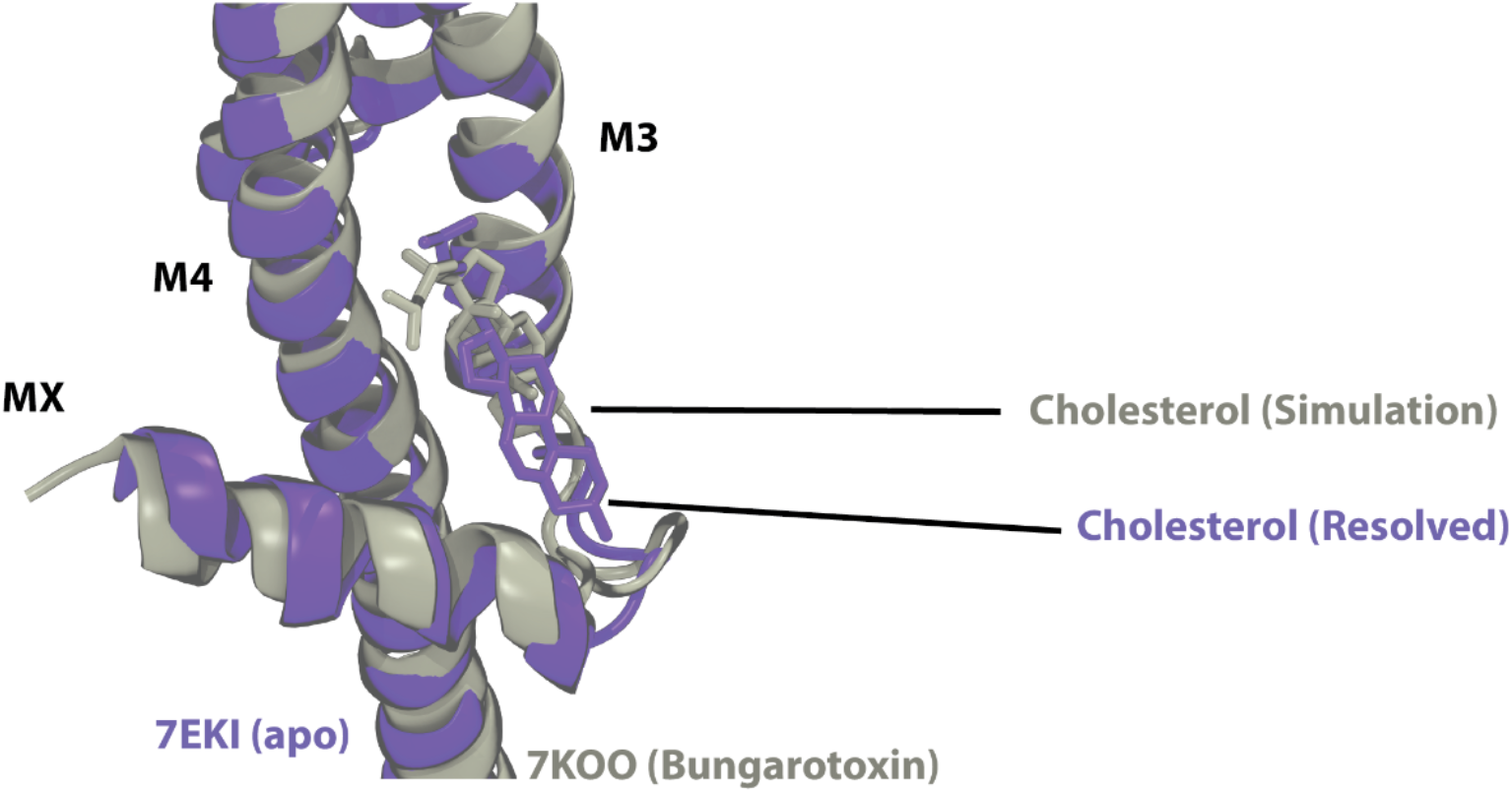
Cholesterol sites. Comparison of the predicted cholesterol binding site in the a-bungarotoxin structural model (gray, PDB: 7KOO) and the resolved cholesterol binding site in the apo structural model (purple, PDB: 7EKI).

**Figure 6S1.**
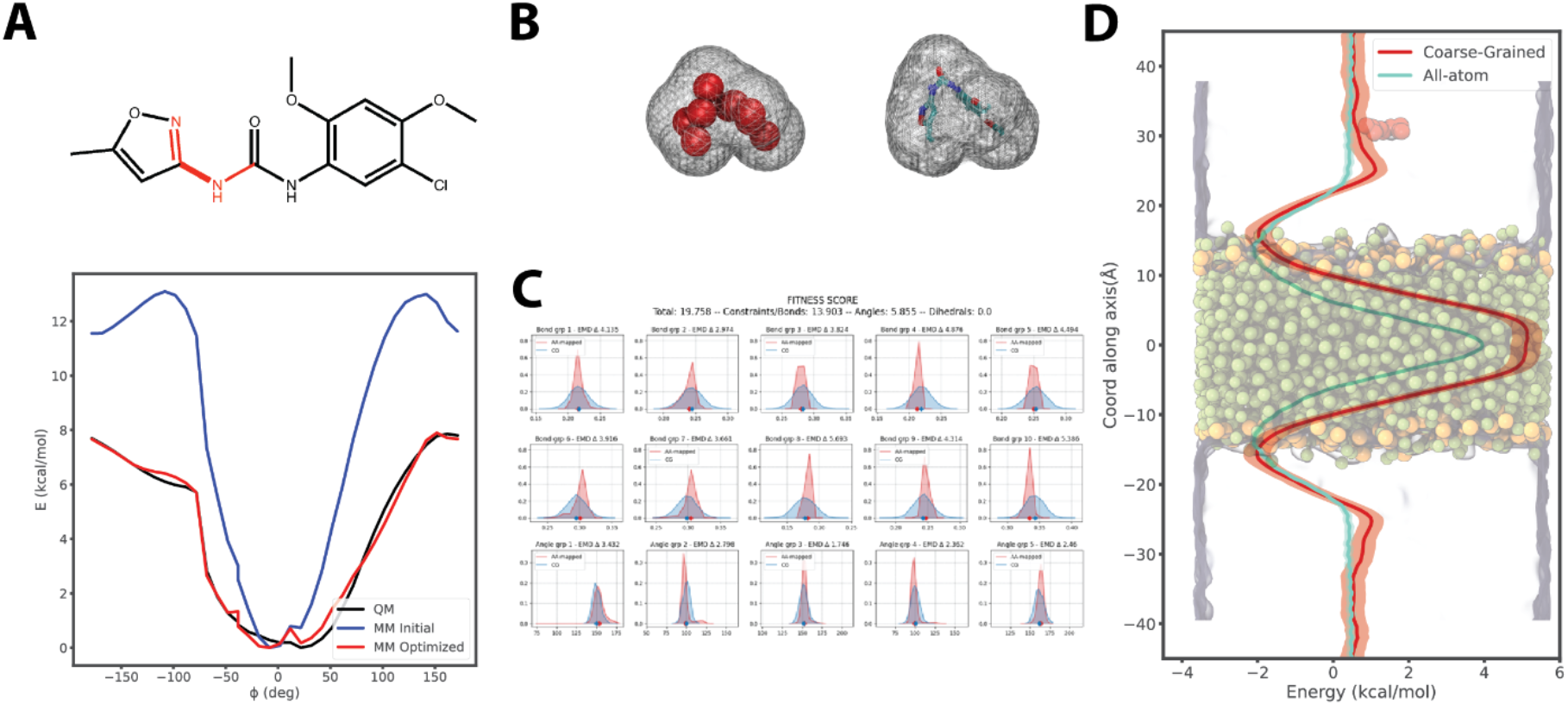
Optimization and validation of PNU parameters. **A**. Optimized energy landscape of one torsion angle parameter of the atomistic PNU. **B**. Comparison of solvent accessible surface area (SASA) between atomistic and coarse-grained PNU. **C**. Bonded term optimization and scoring by SWARM-CG (Empereur-Mot et al., 2020) for PNU. **D**. Comparison of bilayer (POPC) permeation free energy of atomistic and coarse-grained PNU.

**Figure 6S2.**
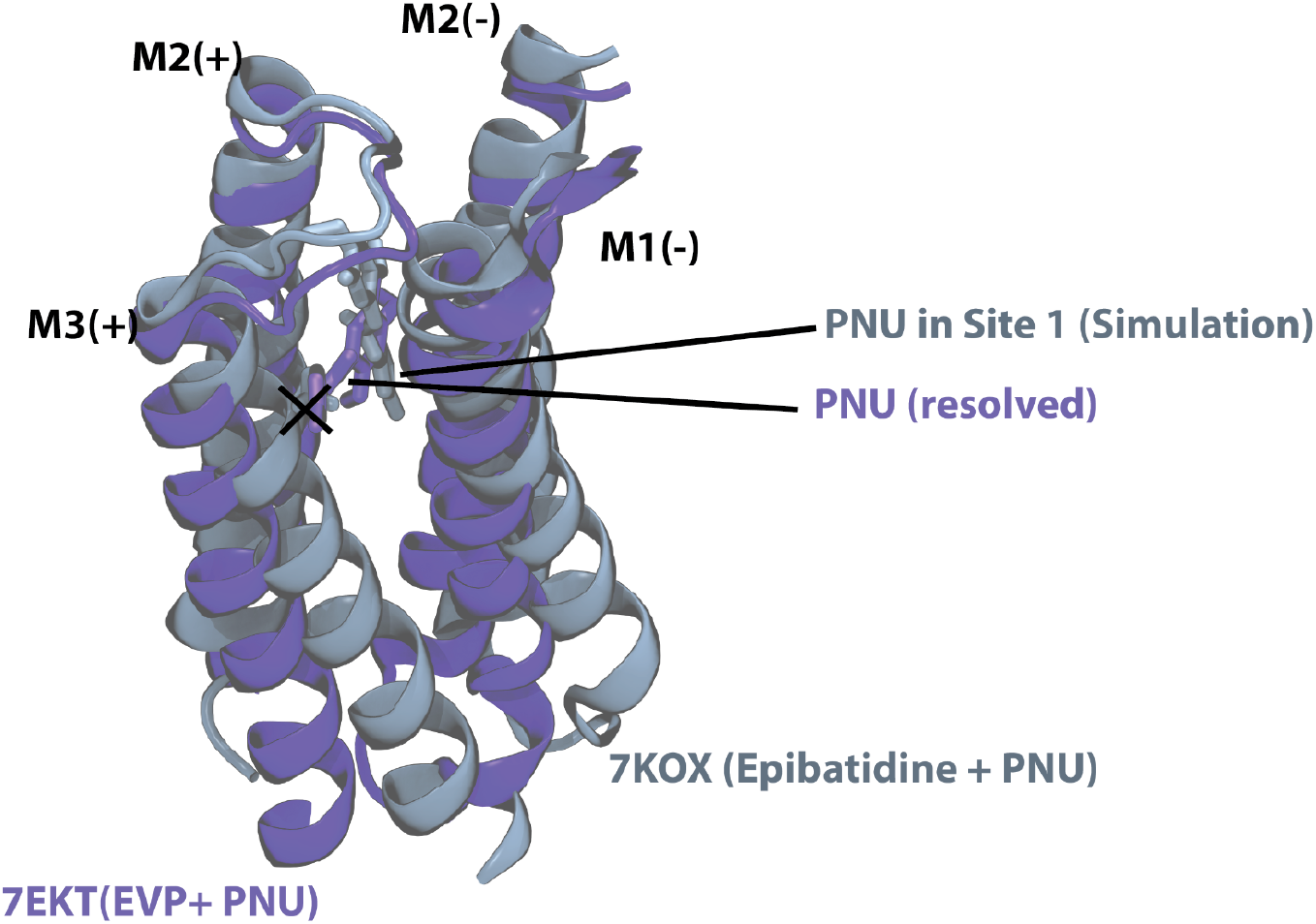
PNU sites. Comparison of predicted PNU binding site in Epibatidine+PNU structural model (gray, PDB: 7KOX) and resolved PNU binding site in EVP+PNU structural model (purple, PDB: 7EKT). The predicted PNU bound with its long axis parallel to the membrane plane while the experimentally resolved PNU had this axis oriented perpendicularly.

**Figure 6S3.**
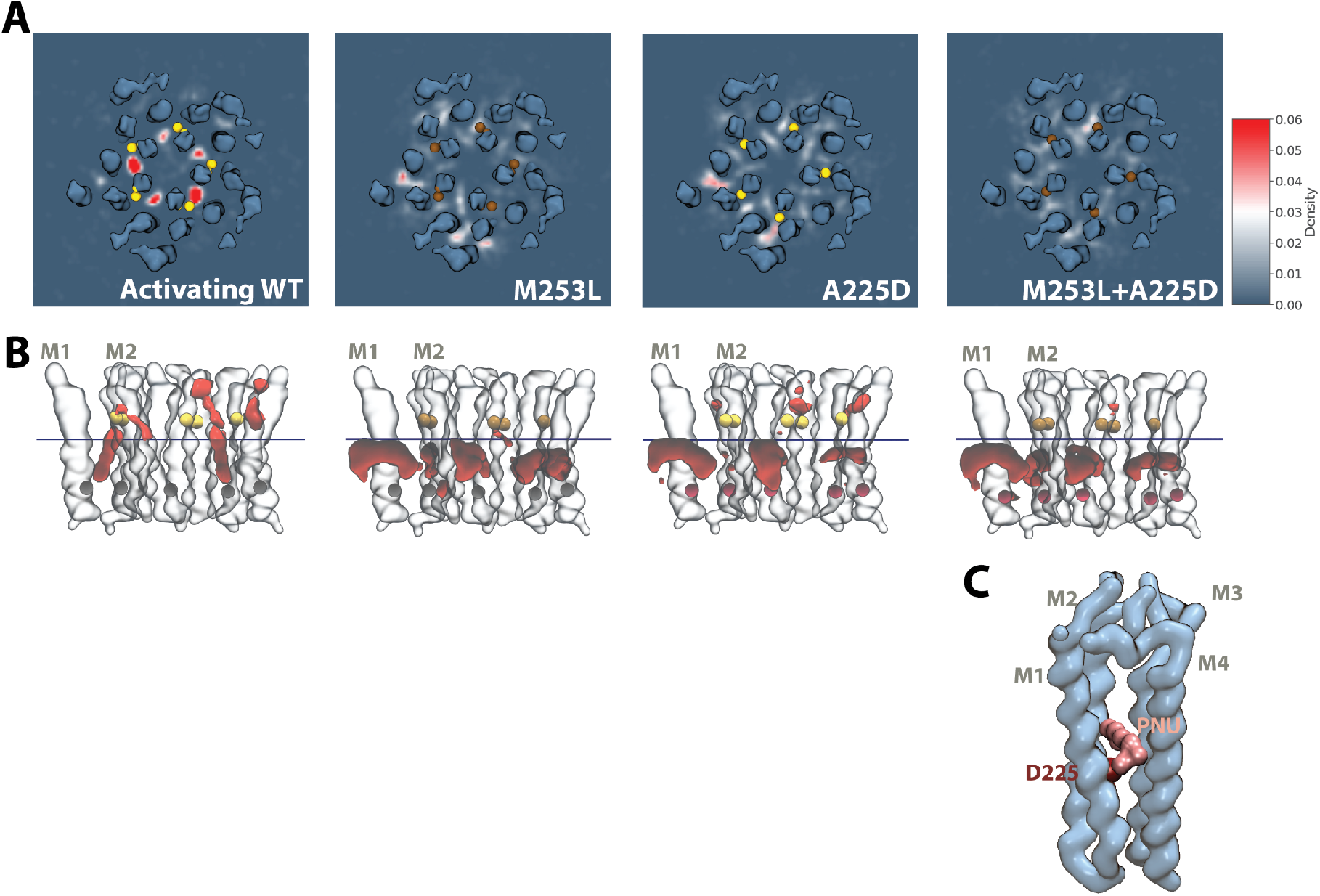
Simulations of mutant systems confirm the PNU binding sites. **A**.The PNU density map derived from 20 * 4 μs simulations in different mutant systems. **B**. The corresponding PNU density (red) in different mutant systems. The slice shown in A-D was plotted as a blue horizontal line. The residue 253 was shown as either yellow (M) or brown (L) bead; The residue 225 was shown as either grey (A) or ruby (D) bead. Only M1, M2 were shown as transparent surfaces. **C**. A snapshot from the CG simulations showed the possible binding mode of PNU (pink) in the mutant system including interaction with D225 (dark red).

**Figure 6S4.**
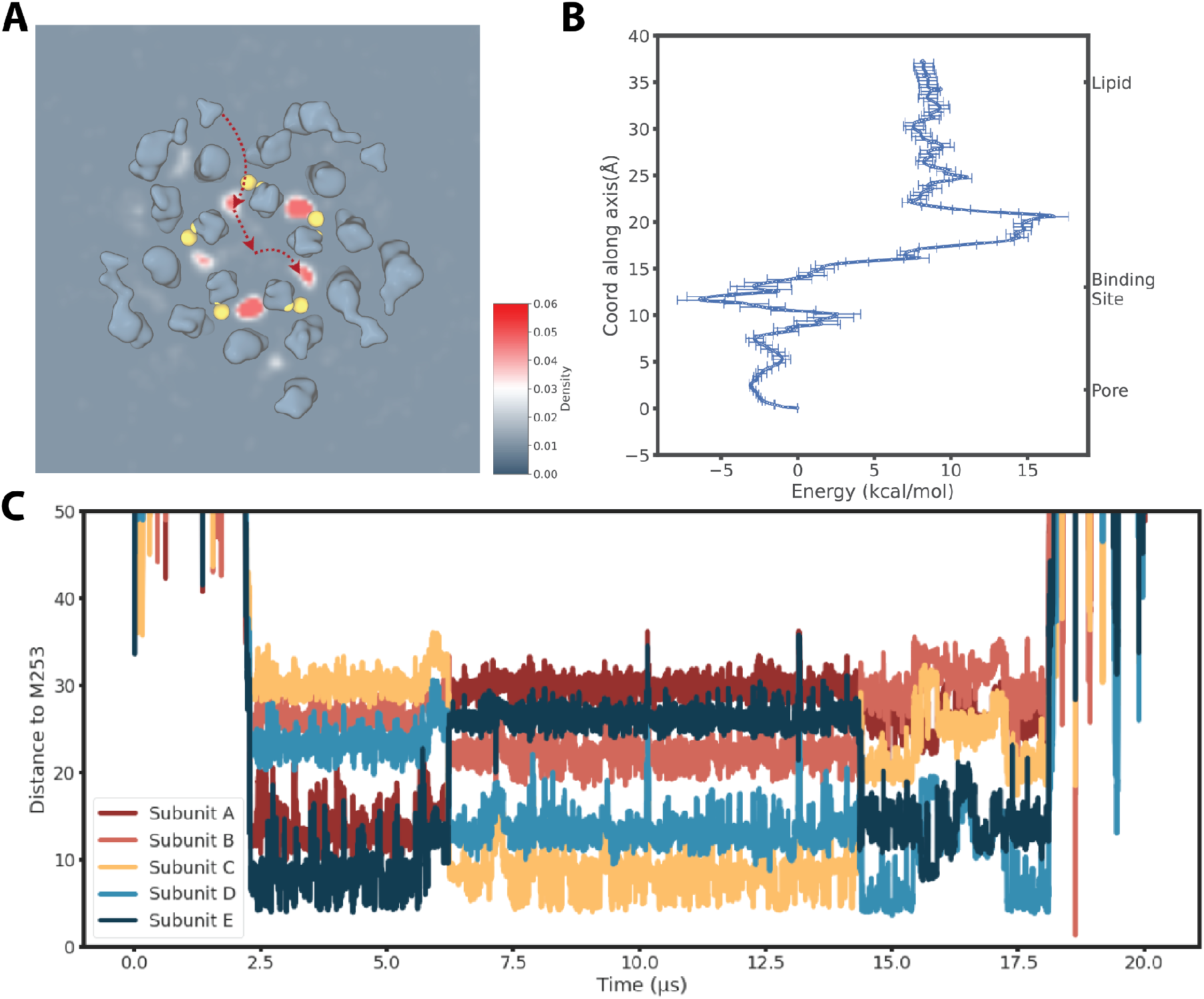
Spontaneous PNU transits between site-1 interfaces in coarse-grained simulations. **A**. Illustration of a PNU molecule diffused from site 1 in one of the subunits across the pore into site 1 located in another subunit in CG simulations.**B**. Free energy landscape of PNU transiting from the pore into the membrane via site 1 by atomistic umbrella sampling simulations. **C**. The time evolution of the distance of one PNU molecule with five M253 in one CG simulation. Aside from entering/exiting the site 1 from/into the bilayer region at ∼2 μs and ∼18 μs. The switching of binding sites can be visualized at ∼6 μs, and ∼14 μs.

**Video 6S5. Movie of spontaneous PNU transition between site-1 interfaces in coarse-grained simulations**.

**Figure 7S1.**
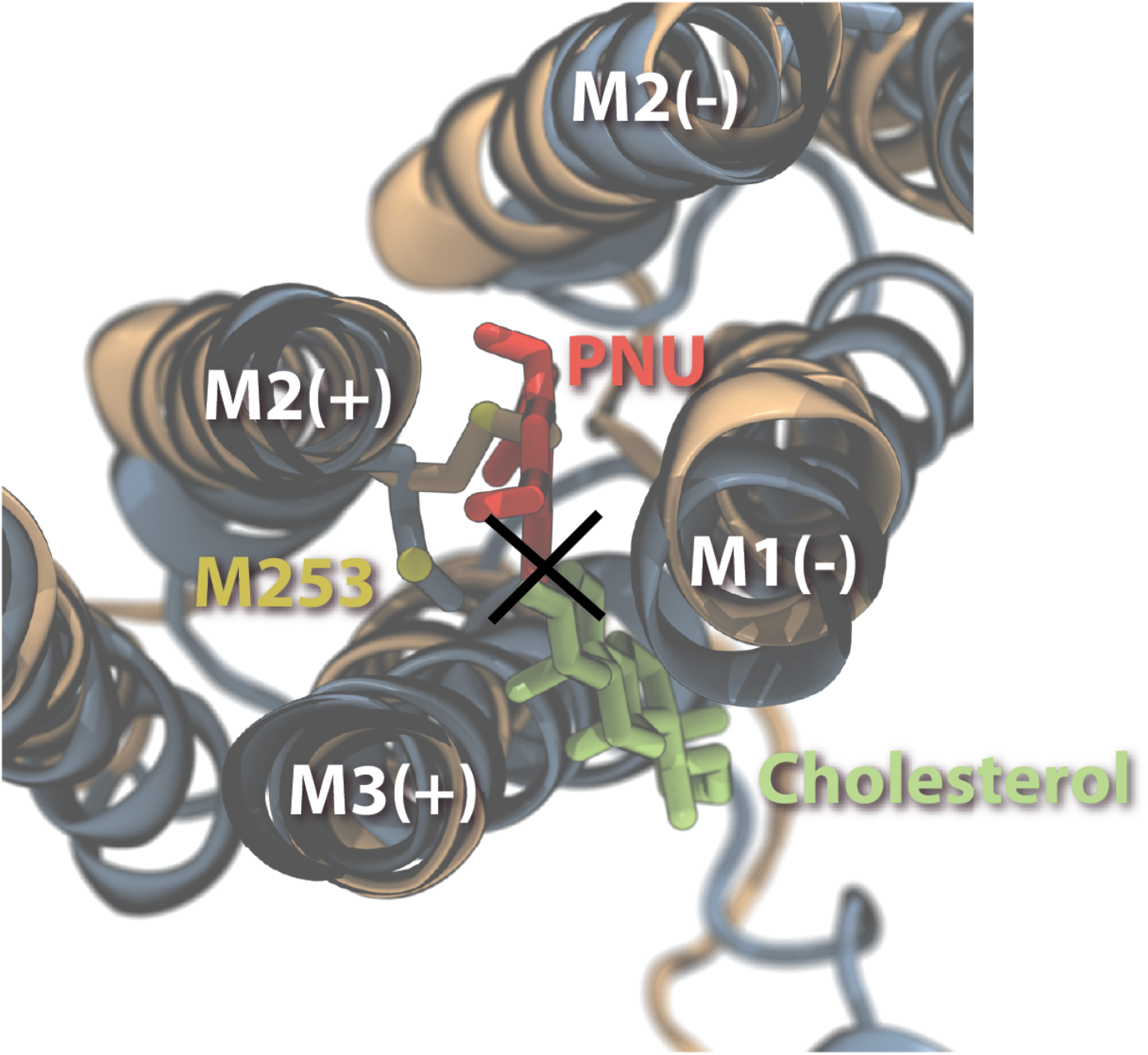
State-dependent interfacial binding. Superposition of the proposed cholesterol binding site in the desensitized state and the proposed PNU binding site in the activated state.

